# Membrane tension regulation is required for wound repair

**DOI:** 10.1101/2024.08.15.608096

**Authors:** Nikita Raj, Martin Weiss, Bart E. Vos, Sarah Weischer, Frauke Brinkmann, Timo Betz, Britta Trappmann, Volker Gerke

**Affiliations:** Institute of Medical Biochemistry, Centre for Molecular Biology of Inflammation (ZMBE), University of Muenster; Münster, Germany; Multiscale Imaging Centre, Cells in Motion Interfaculty Center, University of Münster; Münster, Germany; Bioactive Materials Laboratory, Max Planck Institute for Molecular Biomedicine; Münster, Germany; Third Institute of Physics, University of Göttingen; Göttingen, Germany; Imaging Network, Cells in Motion Interfaculty Centre, University of Münster; Münster, Germany; Cluster of Excellence “Multiscale Bioimaging: From Molecular Machines to Networks of Excitable Cells” (MBExC), University of Göttingen; Göttingen, Germany; Department of Chemistry and Chemical Biology, TU Dortmund University; Dortmund, Germany

**Keywords:** Plasma membrane resealing, Ca^2+^, clathrin-mediated endocytosis

## Abstract

Disruptions of the eukaryotic plasma membrane due to chemical and mechanical challenges are frequent and detrimental, and thus need to be repaired to maintain proper cell function and avoid cell death. However, the cellular mechanisms involved in wound resealing and restoration of homeostasis are diverse and contended. Here, we show that clathrin-mediated endocytosis is induced at later stages of plasma membrane wound repair following the actual resealing of the wound. This compensatory endocytosis occurs near the wound, predominantly at sites of previous early endosome exocytosis which is required in the initial stage of membrane resealing, suggesting a spatio-temporal co-ordination of exo- and endocytosis during wound repair. Using cytoskeletal alterations and modulation of membrane tension and membrane area, we identify membrane tension as a major regulator of the wounding-associated exo- and endocytic events that mediate efficient wound repair. Thus, membrane tension changes are a universal trigger for plasma membrane wound repair modulating the exocytosis of early endosomes required for resealing and subsequent clathrin-mediated endocytosis acting at later stages to restore cell homeostasis and function.

## Introduction

The plasma membrane establishes cellular identity by physically delineating a cell from the extracellular environment. As such, threats to plasma membrane (PM) integrity are frequent, arising from various physiologically experienced mechanical stresses including stretch and fluid flow or external challenges such as pore forming toxins during microbial attack ^[1] [2]^. These recurring challenges often injure the PM resulting in a loss of the phospholipid bilayer integrity and exchange of ions and soluble material between the cell and the extracellular milieu. This uncontrolled exchange eventually leads to loss of cell function and ultimately cell death, and thereby affects tissue integrity at a later stage. Unrepaired PM wounds have been associated with several diseases including muscular dystrophies and ischemic injury, highlighting the importance of maintaining PM integrity in human physiology ^[3]^. To cope with the frequent PM wounds and prevent such pathologies, cells are equipped with mechanisms to efficiently respond to PM injuries, commonly termed plasma membrane repair ^[4]^.

The onset of plasma membrane repair is defined by the entry of extracellular Ca^2+^ through the wound, which triggers the active membrane repair process. The initial phase of the repair process constitutes the immediate establishment of a membrane barrier and the actual PM resealing that terminates the wound-induced Ca^2+^ influx. In the later stage of repair, post-resealing processes aid in restoring homeostasis of the PM to its pre-wounding state. Several cellular mechanisms which are activated by the wound-induced Ca^2+^ influx have been shown to facilitate PM resealing and repair in various cell types ^[5]^. Among others, these can vary from exocytosis-mediated fusion of vesicles ^[6]^, endocytic removal of wounded membrane ^[7]^, ESCRT-regulated membrane shedding ^[8]^, to cytoskeletal rearrangement ^[9]^. The variations are usually attributed to differences in wounding sources, wound sizes, and cell types ^[10]^. However, PM injuries are routinely encountered challenges ^[2]^ and PM repair has been shown to utilize evolutionarily conserved mechanisms already present in the cell (such as exocytosis and endocytosis) ^[11] [12]^. In addition, irrespective of the cell type, the main requirements of PM repair are consistent, ranging from closure of the wound by membrane resealing to restoration of membrane homeostasis after wound repair ^[13]^. Therefore, it remains contended which fundamental mechanisms are relevant to all wound repair processes and how different cellular mechanisms are involved in the central process of plasma membrane repair.

Exocytosis of intracellular vesicles has been recognized as an initial repair response in the resealing of PM wounds in several cells ^[11]^, but the role of these vesicles in membrane repair is disputed, varying from providing membrane to initiating subsequent processes ^[14] [15]^. Endocytosis has also been shown to act in the course of PM resealing by removal of the damaged membrane, thereby establishing a resealed barrier ^[7]^. However, different studies have demonstrated endocytic removal to play a role in other stages of wound repair supporting membrane restructuring ^[16]^. Remodelling of the actin cytoskeleton is yet another mechanism proposed to aid in PM repair by contraction of the wound site for initial resealing ^[17]^, or on the other hand, by acting at later stages of PM repair ^[18]^. Conversely, it has also been reported that disruption of the actin cytoskeleton promotes efficient PM repair, possibly by relieving membrane tension ^[19]^. These contradictory reports raise the question of how and to what extent the various processes that are initiated by the influx of Ca^2+^ upon wounding contribute to both the immediate resealing of the PM bilayer and to the subsequent membrane and cortical remodelling that restores cellular homeostasis.

Recently, we showed that mechanical PM wounding triggers a Ca^2+^-dependent exocytosis of early endosomes that is spatially restricted to the wound site and provides membrane to reseal the injuries ^[20]^. This early endosome exocytosis occurred within seconds of wounding to yield a fast response required to initiate PM resealing and aid in wound closure. Based on these findings, we addressed whether other cellular mechanisms are required for wound repair and restoration of membrane homeostasis. Vascular endothelial cells were chosen as a model system to study PM repair as their membranes are constantly exposed to mechanical challenges due to forces of the circulation rendering them prone to ruptures ^[21]^. Rapid PM wound repair thus has to be initiated to prevent endothelial leakage and inflammatory responses ^[22]^. Here, we show that clathrin-mediated endocytosis is activated near the wound site after early endosome exocytosis and PM resealing have occurred. This compensatory endocytosis is induced at sites of previous early endosome exocytosis suggesting that exocytosis and endocytosis are interlinked events occurring in the course of PM repair. We also show that membrane tension changes caused by cytoskeletal alterations, by varying cell-extracellular matrix tension, by membrane area changes, or by changes in the expression of mechanosensitive proteins, are a central regulator of the repair process that are affected by and control exocytosis and endocytosis. Together, our analysis shows for the first time that exocytic and endocytic processes act cooperatively to regulate membrane tension at various stages of PM repair and thereby collectively facilitate plasma membrane repair in mechanically challenged endothelial cells.

## Results

### Clathrin-mediated endocytosis is activated post plasma membrane resealing in injured HUVEC

We reported earlier that wound repair in mechanically challenged human umbilical vein endothelial cells (HUVEC) requires the Ca^2+^-evoked exocytosis of early endosome (EE), which leads to an accompanying increase in membrane area and thereby resealing ^[20]^. Based on these findings we wondered whether the increase in membrane area is associated with subsequent retrieval of the deposited membrane and therefore, examined the involvement of endocytosis during PM wound repair by analysing different aspects of PM dynamics at various stages of wound repair. We employed two-photon laser ablation to induce site-specific membrane injuries in HUVEC ^[23]^ and first examined the classical clathrin-mediated endocytosis (CME) pathway. In particular, we investigated the dynamics of GFP fusions of the CME-associated proteins Amphiphysin-1 (Amph1) ^[24] [25]^ and Dynamin-2 (Dyn2) ^[26] [27]^ during wound repair. Surprisingly, no significant recruitment of these proteins to the PM, indicative of increased endocytosis, was detected during the course of membrane resealing, which occurs within ∼30 *s* of wounding in HUVEC ^[23]^ (Supplementary Figure S1A, B). This was in contrast to studies reporting an endocytosis-mediated resealing of wounds ^[28]^, but was in line with our observations that EE exocytosis is the major contributor to resealing ^[20]^. Interestingly, at a later stage of repair, Amph1 and Dyn2 formed punctae-like structures in a wave-pattern around the wound site (Figure 1A, B and Supplementary Movies S1 and S2). These structures were observed after resealing of the PM wound, i.e., after 30 – 40 *s* of wounding, and continued for minutes after the wound was closed, occurring in a zone around the wound site (assessed based on FM4-64 wounding dye accumulation marking the resealing site) (Supplementary Figure S1C, D). This indicated that CME was activated near the wound site post wound resealing in HUVEC (also see Supplementary Figure S1E, F). To quantify the CME accumulation observed around the wound site, we implemented a concentric-circle region of interest (ROI) approach ^[20]^ where the distribution of the endocytic punctae across various regions of the cell was analysed with respect to time (see Supplementary Figure S1G). This revealed a striking formation of CME (Amph1 and Dyn2 positive) punctae after wound resealing, predominantly in regions of the cell close to the wound site (Figure 1C, D). No such upregulated endocytic punctae formation was observed in non-wounded cells, pointing towards a wounding-induced response of clathrin-mediated endocytosis and not steady-state endocytosis which occurs always (Supplementary Figure S1H, I).

**Figure 1.**
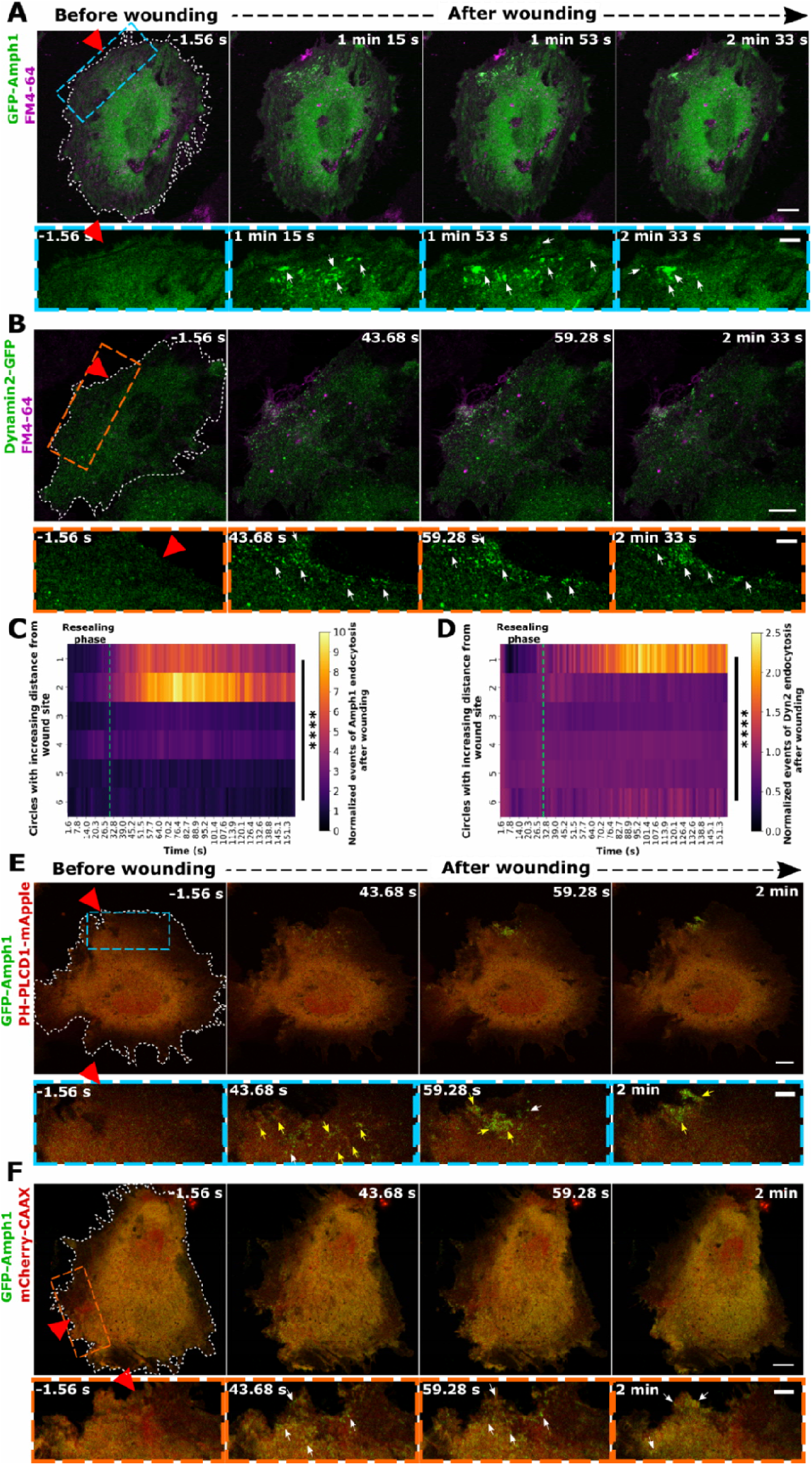
Clathrin-mediated endocytosis occurs near membrane wound sites after resealing. (**A and B**) HUVEC transfected with EGFP-Amphiphysin-1 (green; labelled as GFP-Amph1) (**A**) or Dynamin-2-EGFP (green; labelled as Dynamin2-GFP) (**B**) were kept in buffer containing the membrane-impermeable wounding FM4-64 (magenta) and laser injured at t = 0 *s* during time-lapse microscopy. Representative time-lapse still images before (leftmost panel) and after wound resealing are shown here (resealing typically occurs within 30 *s* post wounding in HUVEC (*22*)). Boxed areas below show higher magnifications of the regions around the wound site at each time point for EGFP-Amphiphysin-1 (blue box) and Dynamin-2-EGFP (orange box). White arrows depict sites of EGFP-Amphiphysin-1 and Dynamin-2-EGFP recruitment indicative of clathrin-mediated endocytosis (CME). (**C and D**) Quantification of the increase in CME events marked by EGFP-Amphiphysin-1 (**C**) and Dynamin-2-EGFP (**D**) punctae, following wounding as a function of time and increasing distances from the wound site (see Supplementary Figure S1G), represented as heatmaps. The green dotted line indicates the time point of resealing after wounding as revealed by cessation of FM4-64 influx (referred to as ‘resealing phase’). The punctae count of endocytic events is normalized to the baseline endocytic count before wounding in each cell and to the area of each circle ROI. (**E and F**) Dynamics of EGFP-Amphiphysin-1 (green) in HUVEC co-expressing the PIP2 lipid marker, PHPLCD1-mApple (**E**) or the general membrane marker, mCherry-CAAX (**F**) (both displayed in red) was recorded after laser ablation and still images are shown here. Boxed areas around the wound site are magnified below for each time point. White arrows indicate sites of Amph1-positive CME and yellow arrows indicate CME events colocalizing with the corresponding lipid markers. Red triangles indicate the sites of laser injury and wounded cells are outlined in white dashes. Scale bars, 10 μm; for zooms, 5 μm. *n* = 18 – 25 cells for (C) and (D), from 3 independent experiments, with means shown here. Statistical comparisons were performed with one-way ANOVA with Kruskal Wallis test for (C) and (D). \*\*\*\**P* < 0.0001.

Next, we assessed whether other pathways of endocytosis are also activated post wounding in HUVEC. Owing to the abundance of caveolar endocytosis in endothelial cells ^[29]^, we first analysed the dynamics of caveolin-1 following PM wounding. However, no change in the localization was observed around the wound site, also at later stages of wound repair (Supplementary Figure S2A, B). Similarly, markers for other clathrin-independent endocytic pathways such as, the CG (CLIC/GEEC for clathrin-independent carrier/GPI-anchored protein-enriched early endosomal compartment) pathway mediated by Cdc42 ^[30]^, RhoA-dependent endocytosis ^[31]^, and the Arf6-dependent pathway ^[32]^, showed no endocytic punctae formation during the course of wound repair (Supplementary Figure S2C - E), suggestive of a specific activation of CME post wounding in HUVEC.

Extracellular calcium influx is a key trigger for initiating wounding responses including the immediate Ca^2+^-triggered EE exocytosis, and is crucial for membrane resealing and associated repair ^[15]^. To confirm that the CME punctae formed are indeed wounding responses following membrane resealing, Ca^2+^ was excluded from the extracellular environment during wounding to prevent membrane addition by Ca^2+^-triggered EE exocytosis. This caused a defect in membrane resealing as observed previously ^[33] [20]^, and also an inhibition of the upregulation of CME punctae that normally occurred after wound resealing (Supplementary Figure S3). As intracellular Ca^2+^ concentrations return to low resting levels upon resealing ^[20]^, high intracellular Ca^2+^ is not required in the later stages of wound repair indicating that post-resealing endocytosis is not driven by Ca^2+^.

Classical CME events are initiated at PM sites enriched in phosphatidylinositol-4,5-bisphosphate (PIP_2_) which can recruit several endocytic factors ^[34]^. To probe whether the endocytic punctae formed upon wounding correspond to CME events at the membrane, the association of Amph1 with the PIP_2_ marker, PH-PLCD1-mApple was investigated. Punctate-like structures of PH-PLCD1-mApple were observed immediately following wounding and after resealing, as noted before in HUVEC ^[18]^, indicating accumulations of PIP_2_ rich membrane areas, which are potential sites for CME. Moreover, the Amph1 punctae formed after resealing colocalized with the PIP_2_ punctae around the wound site with similar kinetics ^[35]^ (Figure 1E). To rule out an association of the Amph1 punctae with stochastic PM structures formed due to wounding or ruffling of the membrane, the dynamics of the general PM marker, mCherry-CAAX was recorded. Hardly any mCherry-CAAX punctae were observed following wounding ^[18]^ and furthermore, no colocalization was seen for Amph1 punctae with the weak CAAX-positive structures (Figure 1F). Together, this indicates that clathrin-mediated endocytosis events occur close to the wound site after membrane resealing in HUVEC, suggestive of a role of endocytosis in the later stages of membrane repair.

### Clathrin-mediated endocytosis occurs at sites of previous early endosomal exocytosis in the course of membrane wound repair

In the initial stages of wound repair, EE exocytosis events occur within the first few seconds of PM damage to promote resealing ^[20]^. This immediate Ca^2+^-evoked exocytosis of EE results in a clustered PM appearance of the early endosomal protein, transferrin receptor (TfR) near the wound site, highlighting abundant membrane deposition. Such membrane deposition at and around the wound site likely results in excess membrane material and we therefore tested whether the initiation of CME in the later stages of wound repair, i.e. after the resealing phase, is a compensatory response to the abundant EE exocytic membrane depositions. To analyse this potential association, we simultaneously recorded the dynamics of a pH-sensitive fluorescent EE reporter, TfR-pHuji, and GFP-Amph1 during wounding. Here, the TfR-pHuji marks EE exocytotic fusion events due to the neutralization-induced increase of the pHuji fluorescence occurring upon fusion of endosomes with the PM. The analysis revealed a striking spatial overlap between the exo- and endocytic events near the wound site (Figure 2A and Supplementary Movie S3). As expected, the GFP-Amph1 punctae appeared much later in time as compared to the immediate TfR-pHuji exocytosis events. However, the Amph1 punctae formed around the same sites near the wound where the TfR-pHuji positive endosomes had previously fused with the PM. Likewise, a spatial association was also seen for Dyn2 endocytic punctae with TfR-pHuji clusters that had appeared earlier after wounding (Figure 2B and Supplementary Movie S4). The spatial proximity between EE exocytosis and CME occurring at different time points after wounding was next evaluated using a nearest-neighbour analysis where the distance between the CME events to the previous EE exocytic sites was measured over time (see Supplementary Figure S4A and Supplementary Movie S5). The analysis detected a wounding-induced spatial association of CME to exocytotic TfR clusters which occurred after resealing near the wound site, but not in distant parts of the cell (Figure 2C, D and Supplementary Figure 4B, C). Such spatial proximity suggests a correlation of EE exocytosis events occurring during resealing and the subsequent internalization of membrane via CME at the same sites. The enrichment of PIP_2_ near the wound site during resealing observed earlier ^[18]^, furthermore indicates that these PIP_2_ accumulations act as potential ‘hotspots’ to define sites, which initiate CME occurring later, suggesting that EE exocytosis and CME are accompanying processes during membrane repair.

**Figure 2.**
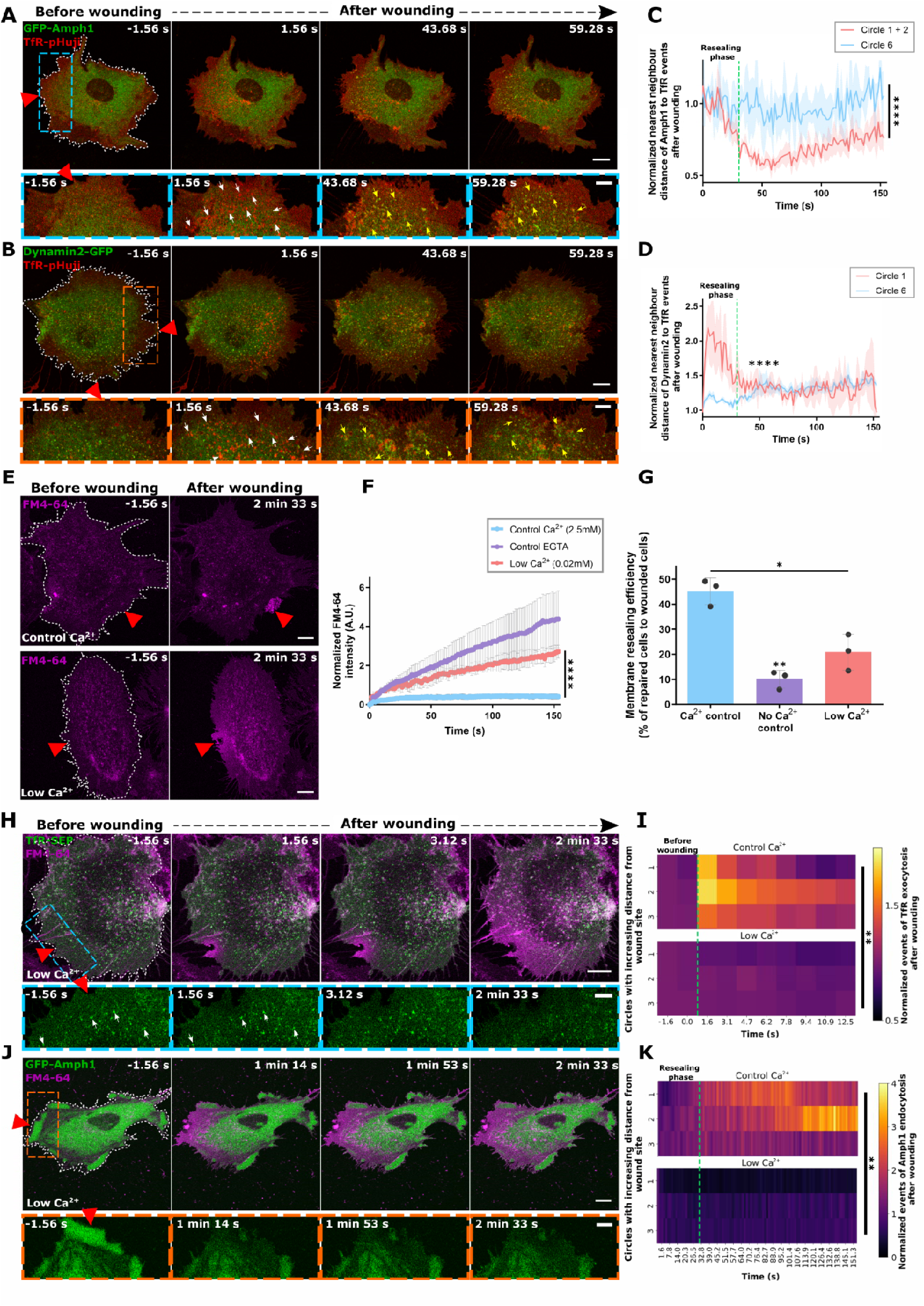
Clathrin-mediated endocytosis which occurs post resealing is associated with sites of previous early endosome exocytosis. (**A and B**) HUVEC co-expressing TfR-pHuji (red) and EGFP-Amphiphysin-1 (**A**) or Dynamin-2-EGFP (**B**) (both shown in green) were laser wounded to assay for spatial association and sequential still images are shown. Higher magnifications of an area around the wound site are shown below for each time point (blue box and orange box for TfR-pHuji with EGFP-Amphiphysin-1 or with Dynamin-2-EGFP, respectively). White arrows indicate sites of TfR-pHuji positive EE exocytosis events near the wound site (increase of pHuji fluorescence due to pH neutralization after exocytosis) and yellow arrows indicate clathrin-mediated endocytosis occurring at sites which coincide with previous wounding-induced EE exocytosis events. **(C and D**) Quantification of the spatial association between wounding-induced early endosomal exocytosis near the wound site (TfR-pHuji signal in circle 1) and subsequent clathrin-mediated endocytosis (EGFP-Amphiphysin-1 punctae for (C) and Dynamin-2-EGFP for (D)) in the cell, was performed by a nearest-neighbour approach (see Supplementary Figure S4A). Spatial proximity between the endocytic proteins and the EE exocytosis events was plotted after normalization to the distance before wounding, for regions close to the wound site (circles 1 and 2 or circle 1 alone, red line) as well as a region far away from the wound site (circle 6, blue line). The green dotted line on the graph indicates the time point of resealing which is approximately 30 *s* after wounding (‘resealing phase’). The dip in proximity observed after resealing is indicative of endocytosis occurring at or close to sites of previous EE exocytosis. (**E**) HUVEC were subjected to laser wounding in the presence of high (2.5 mM, upper panels) or low extracellular Ca^2+^ (0.02 mM, lower panels) and the wounding dye, FM4-64 (magenta). Representative images taken before and after wounding are shown here, indicating impaired membrane resealing at low extracellular Ca^2+^ (continuous influx of FM4-64 over the entire cell). (**F and G**) Quantification of membrane resealing of cells kept in low Ca^2+^ as compared to cells in normal extracellular Ca^2+^ (control Ca^2+^) and cells in EGTA, i.e. in the absence of extracellular Ca^2+^ (control EGTA). Resealing kinetics after laser injury as revealed by FM4-64 uptake are shown in (**F)** and results of mechanical scrape injury assay revealing population level efficiencies (represented as percentages) are depicted in (**G)**. **(H and J**) Still images of HUVEC expressing TfR-SEP (**H**) or EGFP-Amphiphysin-1 (**J**) (both in green) laser injury in the presence of low Ca^2+^ and FM4-64 (magenta) are displayed. Higher magnifications of the boxed areas are shown below for each time point and white arrows in TfR-SEP indicate the punctae which were already present before wounding. (**I and K**) Quantification of wounding-induced TfR-positive EE exocytosis (**I**) and Amph1-labelled endocytosis (**K**) in the presence of low Ca^2+^ (bottom panels) or control Ca^2+^ (2.5 mM Ca^2+^, top panels), are shown as heatmaps for regions close to the wound site (circles 1-3, also see Supplementary Figure S4D, E). Data is normalized to the initial punctae count for each fusion protein prior to wounding as well as the area of the circle ROI. The green dotted line indicates the time point of wounding for (I) (referred to as ‘before wounding’) and the time point of resealing for (K) (‘resealing phase’). Note the lack of EE exocytosis as well as CME events after wounding in the presence of low Ca^2+^ as compared to the controls. Red triangles highlight wound sites and white dashes indicate the wounded cells. Scale bars, 10 μm; for zooms, 5 μm. Mean ± SEM plotted with *n* = 18 - 20 cells for (C) and (D), mean ± SD plotted for (F) and (G, with distribution) with *n* = 24 – 31 cells (F), and means plotted for (I) and (K) with *n* = 24 – 31 cells. Data are pooled from 3-4 independent experiments. The statistical tests used for comparison were: two-tailed Mann-Whitney U test for (C) and (D), one-way ANOVA with Kruskal–Wallis test (F), repeated measures one-way ANOVA with Tukey’s test for (G) and two-way ANOVA with Tukey’s test for (I) and (K). * P < 0.05; **P < 0.01; ****P < 0.0001.

To further analyse the association between EE exocytosis and CME during wound repair, we individually blocked these two processes. First, we interfered with EE exocytosis occurring during the resealing stage. As an influx of high extracellular Ca^2+^ level is required for this EE exocytosis in HUVEC ^[33] [20]^, we employed wounding in low extracellular Ca^2+^ (0.02 mM) as a tool to block EE exocytosis and membrane resealing and thus study the association between EE exocytosis and CME. Such condition resulted in a continuous influx of the membrane-impermeable wounding dye, FM4-64 following PM wounding by laser ablation, confirming a defect in membrane resealing (Figure 2E, F). In addition, we also used a population wounding assay – mechanical scrape injury – to verify the inhibition of wound resealing in low Ca^2+^ conditions (Figure 2G). As expected, low extracellular Ca^2+^ led to an inhibition of wounding-induced EE exocytosis as revealed by the lack of appearance of brightly fluorescent punctae containing the pH-sensitive fluorescent EE reporter, TfR-SEP (super ecliptic pHluorin) (Figure 2H, I and Supplementary Figure S4D). To study CME in wound repair, here and in later analyses we focused on Amph1 since Dyn2 is also involved in various clathrin-independent pathways ^[31]^. Intriguingly, wounding in the presence of low Ca^2+^ in addition to preventing EE exocytosis events also resulted in a strong obstruction of Amph1 positive CME (Figures 2J, K, and Supplementary Figure S4E), as seen in Ca^2+^-free media as well (Supplementary Figure S3). These results suggested that the upregulation of CME events that occurs at later stages of repair is dependent on membrane resealing and therefore, the EE exocytosis events which promote resealing.

We next examined the interdependence of wounding-induced EE exocytosis and CME by blocking the other associated process, clathrin-mediated endocytosis. We first employed Dynasore, a well-established inhibitor of CME (Supplementary Figure S5A) which blocks Dyn2 function ^[36]^. This resulted in a major resealing defect in HUVEC (Figure 3A, B). To verify that the membrane resealing defect is due to blocking CME (as Dyn2 is involved in various endocytic pathways ^[31]^), we utilized another CME inhibitor, Pitstop-2, which blocks interactions of CME proteins with the clathrin heavy chain ^[37]^ (Supplementary Figure S5B). Wounding of Pitstop-2 treated HUVEC revealed a marked membrane resealing defect as well (Figure 3C, D). However, the long-term block of CME by Dynasore and Pitstop-2 in these experiments affects steady-state endocytosis and therefore depletes the pool of TfR positive EE already prior to wounding (see Supplementary Figure S5A, B) ^[38]^. Therefore, we speculated that such depletion of EE precludes any wounding induced EE exocytosis required for resealing and thereby causes the observed defects in membrane resealing ^[38]^. Indeed, Dynasore-treated HUVEC showed a marked reduction of wounding-induced EE exocytosis events (Figure 3E, G and Supplementary Figure S5C), along with an abrogation of Amph-1 dependent CME (Figure 3F, H and Supplementary Figure S5D). These results indicated that blocking CME in the long-term affects PM repair by depletion of EE and thus inhibition of any EE exocytosis required for resealing and that steady-state CME is required to replenish the pool of early endosomes required for rapid responses to membrane wounding. To rule out any off-target effects of the pharmacological endocytic inhibitors, we also depleted the clathrin heavy chain (siCLTC) by siRNA mediated knockdown to specifically inhibit steady-state CME, as verified by the restricted transferrin cargo uptake observed in siCLTC cells (Supplementary Figure S5E, F). Laser injury of HUVEC after clathrin heavy chain knockdown also demonstrated a major PM resealing defect, confirming the requirement of steady-state CME and the resulting presence of an abundant EE pool for efficient PM repair (Figure 3I, J). Corroborating the laser wounding results, population-based scrape injury assays of Dynasore treated and siCLTC knockdown cells also showed impairments in HUVEC PM resealing (Figure 3K, L). Finally, we substantiated the specific involvement of CME in the course of wound repair by examining the functional role of clathrin-independent pathways during wounding. In line with the lack of an association of clathrin-independent endocytic pathways with the wound site, a functional involvement of these pathways in membrane resealing was also not observed (Supplementary Figure S5G, H). Together, EE exocytosis and its associated compensatory CME are indispensable for the various stages of PM repair process in HUVEC, most likely by modulating membrane area to promote resealing and later membrane remodeling.

**Figure 3.**
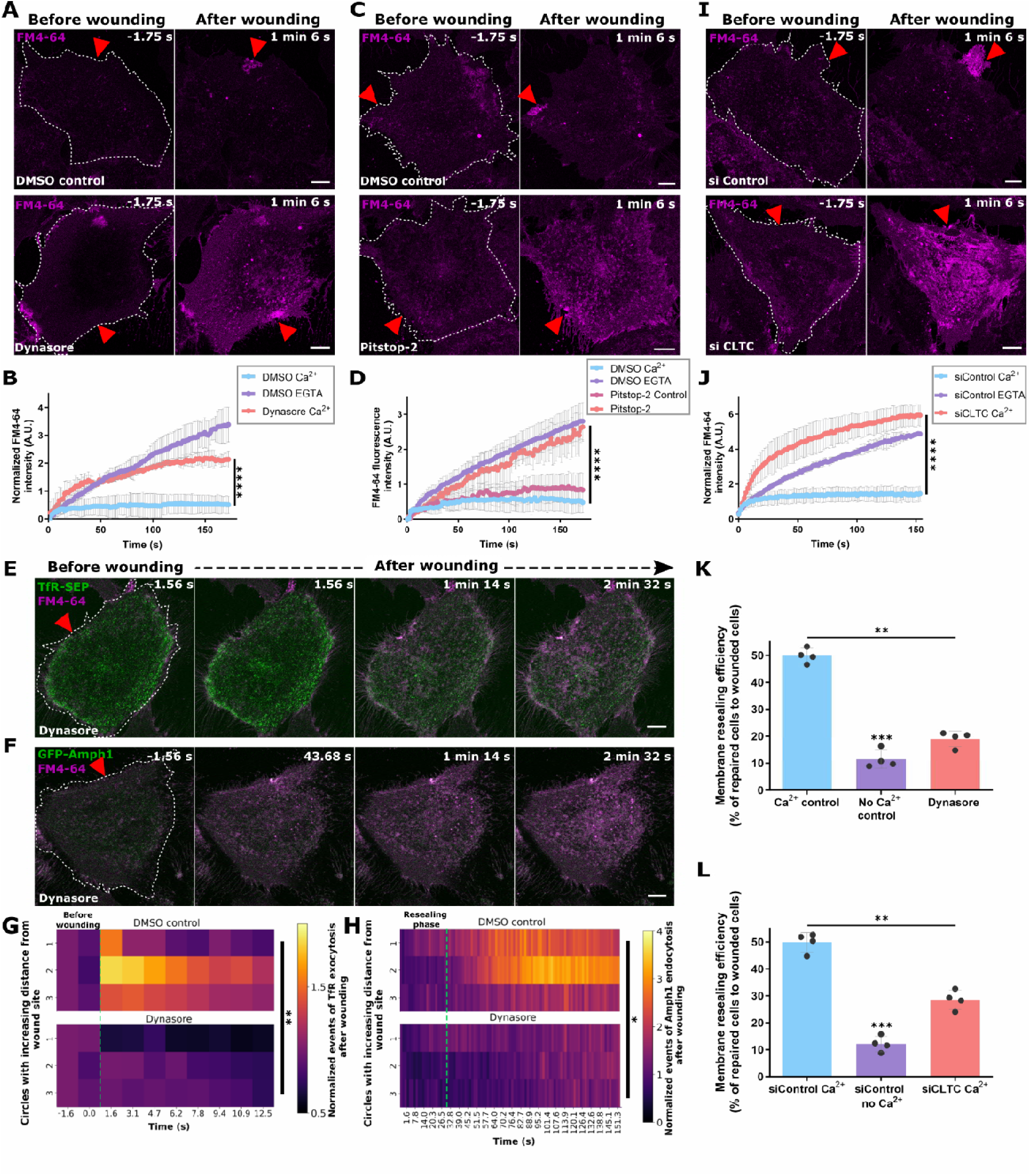
Inhibition of clathrin-mediated endocytosis interferes with wounding-induced EE exocytosis and membrane repair. **(A and B)** Wounding of HUVEC treated with the dynamin inhibitor, Dynasore (bottom panel) in the presence of FM4-64 (magenta) shows membrane resealing defects as opposed to DMSO control (top panel). Representative images shown in (**A**) and quantification of resealing kinetics displayed in (**B**). **(C and D)** HUVEC treated with the CME inhibitor, Pitstop-2 or controls - DMSO or Pitstop-2 control, were laser wounded in the presence of FM4-64. Representative fluorescence images shown in (**C**) as well as quantification of membrane resealing in (**D**) indicate impaired membrane repair. **(E and F)** HUVEC expressing TfR-SEP (**E**) or EGFP-Amphiphysin-1 (**F**) were treated with Dynasore and laser injured in the presence of FM4-64. Images taken immediately after wounding are displayed to reveal EE exocytosis (**E**) and after resealing to detect CME (**F**). Note that both, wounding-induced EE exocytosis and post-resealing CME are impaired following long term Dynasore treatment. **(G and H)** Quantification of the effect of Dynasore on wounding-induced TfR-positive EE exocytosis events (**G**) or post-resealing Amph1-positive CME events (**H**), occurring around the wound site (circles 1-3) as heatmaps. Events are normalized to the frame before wounding and circle areas. The green dotted line marks t = 0 s for wounding in (**G**) and t = 30 s for resealing in (**H**). **(I and J)** HUVEC transfected with siControl or siRNA directed against clathrin heavy chain (siCLTC) were subjected to laser wounding in the presence of FM4-64 (magenta) (**I**) and quantification of resealing kinetics (**J)** are shown. **(K and L)** Resealing efficiencies of cells treated with Dynasore (**K**) or siCLTC (**L**) after mechanical scrape injury as compared to controls. Data are represented as a percentage. Red triangle indicates wound ROI, wounded cells are outlined in white dashes, and scale bars = 10 μm. Mean ± SEM plotted for (D), mean ± SD plotted for (B), (J), (K) and (L), and means plotted for (G) and (H). *n* = 18 – 24 cells for (B, D, J, G and H). All data were pooled from 3-4 independent experiments. The following statistical tests were used for comparison: one-way ANOVA with Kruskal-Wallis test for (B, D, and J), two-way ANOVA with Tukey’s test (G and H), and repeated measures one-way ANOVA (K and L). \**P* < 0.05; \*\**P* < 0.01; \*\*\**P* < 0.001; \*\*\*\**P* < 0.0001.

### Wound-induced EE exocytosis and subsequent clathrin-mediated endocytosis events are linked to membrane tension

The induction of EE exocytosis and CME occurring after wounding at various temporal stages prompted us to study how these events are interlinked during PM repair. Earlier studies have shown that membrane area is regulated by membrane trafficking processes, such as exocytosis and endocytosis activated during PM repair ^[39] [7] [20]^. It was also postulated that changes in membrane area are inversely correlated to changes in membrane tension ^[40] [41]^. Therefore, we inspected the possible role of membrane tension changes in regulating exocytosis and endocytosis events after PM wounding. The apparent membrane tension is considered to depend on both, the in-plane membrane tension and the tension mediated by PM to cortical cytoskeleton attachment ^[42]^. Membrane tension of a cell is therefore affected by altering the actin cytoskeletal machinery, which is necessary for maintaining cellular tension ^[43] [44]^. Treatment with Cytochalasin D (CytoD), an inhibitor of actin polymerization ^[45] [46]^, had been reported to result in lowered membrane tension in HUVEC ^[47]^. Laser injury of HUVEC treated with CytoD showed no impairment in PM repair, rather resealing appeared even slightly enhanced, suggesting a positive role for lowered tension in PM resealing (Figure 4A, B). This result was also validated by mechanical scrape wounding assays (Figure 4C). Inspection of TfR-SEP positive EE exocytosis events (Figure 4D, F, and Supplementary Figure S6A) and Amph1 positive CME (Figure 4E, G, and Supplementary Figure S6B) displayed significantly reduced exocytosis and endocytosis near the wound site of CytoD treated HUVEC. This indicated a possible role of membrane tension in regulating exocytosis and endocytosis during wound repair.

**Figure 4.**
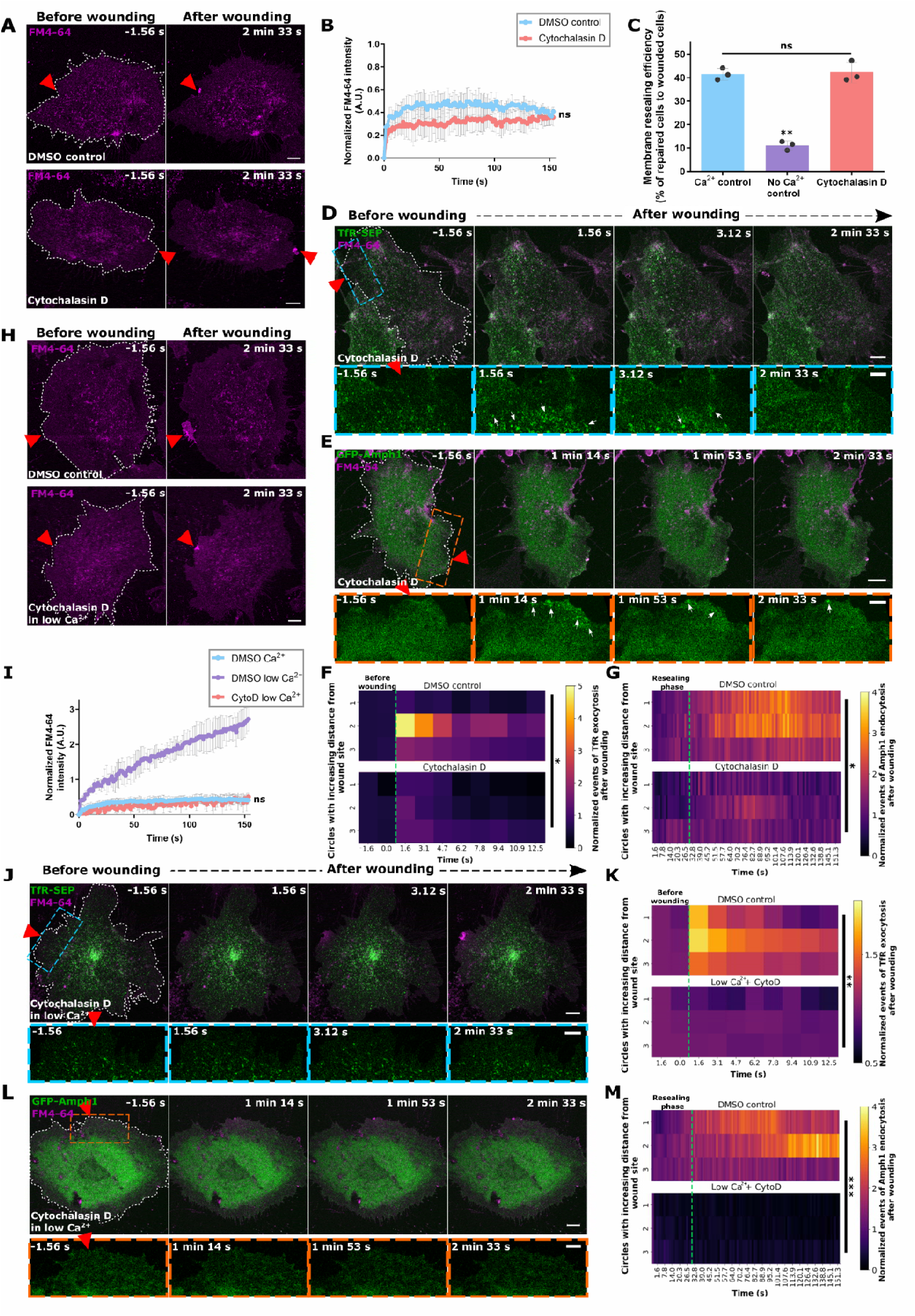
Lowering of membrane tension by blocking actin polymerization promotes membrane resealing. (**A and B**) Still images of laser injury of HUVEC treated with Cytochalasin D (bottom panel) or DMSO control (top panel), in the presence of FM4-64 (magenta) indicate successful resealing (**A**), as also seen in the quantification of resealing kinetics (**B**). (**C**) Resealing efficiency of HUVEC treated with Cytochalasin D or controls and subjected to mechanical scrape injury, represented as a percentage of total wounded cells. **(D and E**) TFR-SEP (**D**) or EGFP-Amphiphysin-1 (**E**) expressing HUVEC (both visible in green) were laser wounded in the presence of Cytochalasin D and FM4-64 and representative stills shown here. Boxed regions around wound sites are magnified below. White arrows indicate EE exocytosis events positive for TfR-SEP (**D**) and clathrin-mediated endocytosis marked by enriched EGFP-Amphiphysin-1 (**E**). **(F and G**) Heatmap representations of wounding-induced punctae count of TfR-SEP (**F**) and EGFP-Amphiphysin-1 (**G**) in DMSO control (top panel) or Cytochalasin D treated cells (bottom panel). Punctae are plotted across time in regions close to the wound site (circles 1-3), and normalized to the initial punctae count and circle area. The green dotted line indicates the time of wounding (**F**) and time of resealing (**G**). (**H and I**) HUVEC were treated with Cytochalasin D or control DMSO and wounded in the presence of low extracellular Ca^2+^ and FM4-64 dye. Representative still images (**H**) and quantification to assess for resealing kinetics (**I**). **(J - M**) HUVEC were transfected with TFR-SEP (**J,** quantified in **K**) or EGFP-Amphiphysin-1 (**L,** quantified in **M**) (both constructs shown in green in the images), followed by Cytochalasin D treatment and laser injury in low extracellular Ca^2+^ and FM4-64 dye (magenta). Boxed areas below each panel show corresponding magnified views around the wound sites. Quantifications of TfR and Amph1 punctae are represented as heatmaps post-wounding and displayed for regions close to the wound site across treatments. Data are plotted as detailed in (F and G). Red triangles, laser injury sites, and white dashes indicate wounded cells. Scale bars, 10 μm; for magnifications, 5 μm. Mean ± SEM plotted for (B), mean ± SD for (C) and (I), and means are plotted for (F), (G), (K) and (M), with *n* = 21 – 31 cells compiled from 3 independent experiments. *P* value was calculated using the following analyses: Wilcoxon paired t-test (B), one-way ANOVA with Kruskal–Wallis test for (C) and (I), and two-way ANOVA with Tukey’s test (F), (G), (K) and (M). *ns*, not significant; * P < 0.05; **P < 0.01; \*\*\**P* < 0.001.

To further discriminate between the involvement of lowered membrane tension or EE exocytosis and subsequent CME during wound repair, low extracellular Ca^2+^ was utilized in the background of CytoD treatment. Interestingly, wounding of HUVEC in the presence of CytoD and low extracellular Ca^2+^ did not inhibit efficient membrane repair. This contrasts the impaired wound repair of HUVEC in low extracellular Ca^2+^ only (Figure 4H, I; as compared to Figure 2E) and points towards a crucial involvement of lowered tension in PM resealing. Moreover, analysis of TfR-SEP and GFP-Amph1 punctae revealed no discernible EE exocytosis and CME events after wounding of HUVEC in low extracellular Ca^2+^ following CytoD treatment (Figure 4J - M and Supplementary Figure S6C, D), even though PM repair was unaffected (as opposed to Figure 2H, J).

A role of lowered membrane tension in facilitating membrane resealing can also be inferred from biophysical calculations. The closure of holes in an examplary phospholipid bilayer is driven by the high energy cost associated with an open hole edge. This cost arises from either the direct interaction between the hydrophobic lipid tails in the membrane and water or from the significant curvature of the lipids. From a physical perspective, this phenomenon can be conceptualized as an enthalpy cost for creating a hole, analogous to classical enthalpy where line tension substitutes for pressure and the hole’s perimeter replaces volume. This leads to an effective enthalpy expression *H_Line_ = E + 2πRλ*, where *E* represents the system’s energy, *R* the radius of the hole, and λ the line tension of the membrane. Conversely, membrane tension can also be expressed in terms of enthalpy. Here, an increase in hole size decreases the overall energy, which is represented by *H_tension_ =E - πR*^2^*σ*, with σ denoting the membrane tension. As a consequence, physical forces drive the system towards a minimum total enthalpy: *H = 2E + 2πRλ - πR^2^σ*. For small values of *R* or hole sizes, the system rapidly reduces *R* to zero, effectively resealing the hole. However, for larger values of *R*, it becomes energetically favorable to increase *R*, as the energy savings from membrane tension become predominant. To determine the critical radius at which the hole will self-reseal, we minimized *H* with respect to *R* by setting 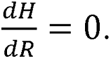 Mathematical calculations reveal that the critical radius of the hole *R_C_* equals *λ/σ* Given that line tension λ is a material property largely invariant during hole closure, it can be considered constant. In contrast, membrane tension σ is a dynamic value affected by the available membrane area. Our results indicate that this tension decreases upon membrane supply by the exocytosis of early endosomes to the wound area ^[20]^. By employing standard values for line tension (λ ≈ 1 pN ^[48]^) and membrane tension (σ≈ 10^-S^Nm^-1 [41]^), the physical principles predict that, in unaltered scenarios, holes in model bilayers of up to 100 nm in diameter will self-heal. Upon local increase of membrane availability (as a consequence of early endosome exocytosis) and inhibition of membrane flow away from the wound (through the dissolution of the actin cortex in the hole-affected area ^[18]^), we can estimate a local reduction in membrane tension, potentially by an order of magnitude or more. This reduction in membrane tension, which is predicted by simple membrane physics, would facilitate efficient and rapid resealing of holes larger than 1 µm (as seen in our studies), suggesting that lowering tension is a robust mechanism for membrane repair. Altogether, our findings and calculations imply that lowered tension is vital for efficient PM resealing and that EE exocytosis and the associated upregulated CME activated in response to wounding possibly aid in regulating membrane tension during wound repair.

### Experimental membrane tension modulation reveals its crucial role in PM repair

As cytoskeletal drugs such as CytoD affect a variety of cellular processes that may lead to off-target effects ^[49]^, we introduced a more precise method for modulating membrane tension by culturing cells on polyacrylamide (PAA) hydrogels of varying substrate stiffnesses (0.2 kPa stiffness selected as ‘soft’ gels and 20 kPa chosen as ‘stiff’ gels here) ^[50]^. Cells grown on such gels have been reported to show specific cytoskeleton/tension variations ^[51]^. To confirm the effect of PAA gel stiffness on the apparent membrane tension of cells grown on them, a membrane tether pulling assay employing optical tweezers was used to measure tether forces in these cells. Here, the measured tether force is correlated to the membrane tension ^[52]^. The corresponding experiments showed that the tether forces are significantly higher in cells grown on the 20 kPa matrix indicating an increased membrane tension in cells cultivated on gels of higher stiffness (Figure 5A).

**Figure 5.**
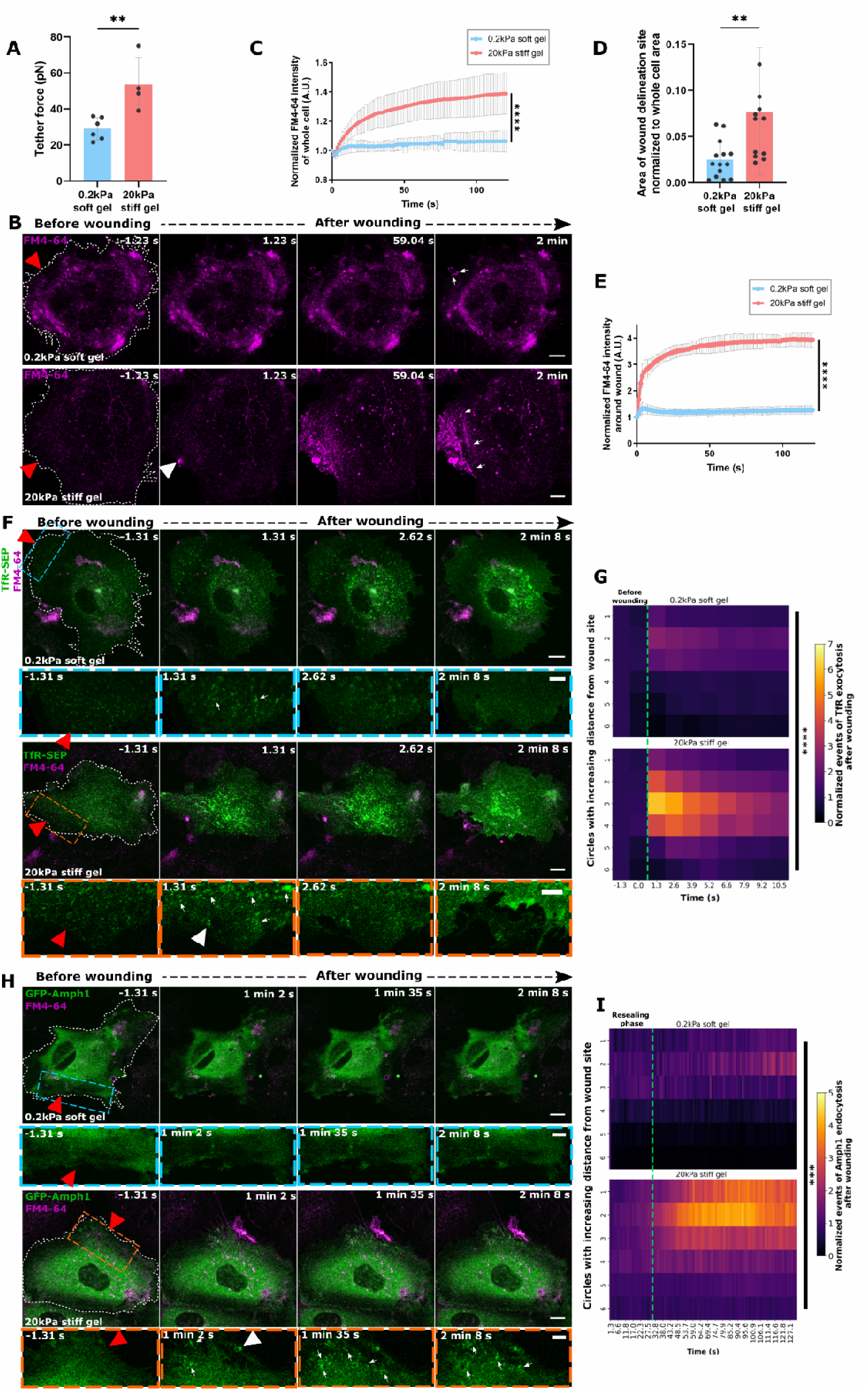
Membrane tension alterations of HUVEC cultured on hydrogels of different stiffnesses affect exo-endocytic processes and wound resealing responses. (**A**) HUVEC grown on collagen-coated polyacrylamide gels of varying stiffnesses of 0.2 kPa (soft gel) and 20 kPa (stiff gel) were subjected to tether pulling experiments to estimate tether force as an indicator of membrane tension. Note the increase in tether force and thus membrane tension in cells grown on the stiffer gels. (**B**) HUVEC grown on soft and stiff hydrogels (as in (A)) were laser injured in the presence of FM4-64 dye (magenta) to analyse resealing capacity. HUVEC grown on soft gels show a stronger basal FM4-64 dye staining due to rounded cell morphology. White arrowhead in the bottom panel (stiff gel) points to FM4-64 accumulation at the wound site observed immediately after wounding which is not seen in wounded cells grown on soft gels. Also note the larger FM4-64 labelled delineation site around the wound in cells grown on stiff gels (highlighted with white arrows labelling the edge of the delineation site in the last time point for both gels). (**C - E**) Membrane resealing efficiency in HUVEC grown on gels of different stiffness was estimated as follows: FM4-64 influx over the entire cell normalized to the signal before wounding (**C**), area of wound delineation site after membrane repair normalized to the whole cell area (**D**), and FM4-64 intensity measurements near the wound in a defined ROI normalized to dye intensity before wounding (**E**). Note the differing resealing efficiencies in cells grown on soft and stiff gels as indicated by the influx of the FM4-64 dye as well as the size of the wound repair sites. (**F**) HUVEC expressing TfR-SEP (green) were grown on soft (top panel) or stiff gels (bottom panel) and laser wounded in the presence of FM4-64. Magnifications of wound sites are given in boxed areas below the images for each stiffness. White arrowhead indicates the TfR-SEP positive EE exocytotic accumulation coinciding with the FM dye accumulation (labelled in the merge panel) at the exact injury site immediately after wounding of a cell grown on stiff gel (bottom panel). White arrows mark the exocytotic TfR-SEP clusters formed after injury. (**G**) Heat map representations of punctae counts of TfR-SEP in cells grown on soft or stiff gels (top and bottom panels, respectively) following wounding as a function of varying distances from the wound site (circles 1-6). The punctae are normalized to initial events before wounding and the corresponding circle area. The green dotted line indicates the time of wounding. Note that EE exocytosis resulting in TfR-SEP punctae is greatly diminished in cells grown on soft gels. (**H and I**) HUVEC cultured on soft or stiff gels were transfected with EGFP-Amphiphysin-1 (green) and images prior to and after wound resealing are shown (**H**). Wound areas are magnified in boxes below. Note the abundance of wounding-induced GFP-Amph1 punctae in the wounded cell grown on stiff gel (white arrowhead indicates the punctae formation at the exact wound site and white arrows indicate the Amph1 punctae around the wound site in the bottom panel), which is not seen in cells grown on soft gels. Quantifications of wounding induced Amph1 punctae in these cells are shown on the right (**I**). Data is plotted as in (G) and the green dotted line indicates the resealing time points here. Red triangle indicates wound ROI, white dashes delineate the wounded cells, and scale bars, 10 μm; for zooms, 5 μm. Mean ± SD shown for (A) with *n* = 4 - 6 cells. Mean ± SEM plotted for (C) and mean ± SD for (D) and (E), from *n* = 33 - 40 cells. Means plotted from 18 - 33 cells for (G) and (I). Data were pooled from 2 – 4 biological experiments. The statistical tests used were: unpaired two-tailed t-test (A), Mann-Whitney U test for (C), (D) and (E), and two-way ANOVA with Tukey’s test for (G) and (I). **P < 0.01; ***P < 0.001; ****P < 0.0001.

We then subjected HUVEC grown on the different hydrogels to laser-ablation wounding assays. It is known that cell spreading increases with increased substrate stiffness ^[53]^ and, as noted before in other cells ^[51] [54]^, HUVEC grown on collagen-coated soft gels showed a more rounded morphology leading to a stronger basal FM4-64 membrane dye staining (Figure 5B, leftmost top panel). Wounding of HUVEC grown on soft gels resulted in unrestricted PM repair, with FM4-64 rich structures restricted to the wound site during repair (Figure 5B and Supplementary Movie S6). In some cases, a strong retraction of the cell was observed during wound repair (possibly owing to the weak adhesion of the cell to the soft gel substrate), nevertheless without affecting the efficient membrane resealing (Supplementary Figure S7A). In stark contrast, injury of HUVEC grown on stiff gels (showing a flatter cell morphology) induced a massive FM4-64 influx indicative of impeded membrane repair (Figure 5B, bottom panel and Supplementary Movie S7A) and in some cases even completely inhibited membrane repair (Supplementary Figure S7B and Supplementary Movie S7B). These results suggested that reduced membrane tension due to growth on soft gels supports PM repair while increased membrane tension experienced in cells grown on stiff gels hindered efficient PM repair. In view of the heterogeneous resealing defects observed in stiff gel-grown HUVEC, membrane resealing was quantified as done previously over the entire cell as well as around the site of wound delineation (Figure 5C – E). All of the analyses revealed a drastic membrane repair defect in stiff gel-grown cells while cells on soft gels resealed efficiently.

Next, we examined the induction of EE exocytosis and CME after wounding in HUVEC grown on soft and stiff gels. TfR-SEP positive clusters revealing EE exocytosis formed immediately upon wounding of cells on both soft and stiff gels; however, to a drastically lower extent in cells grown on soft gels (comparable to the markedly reduced TfR-SEP exocytosis upon wounding of CytoD treated cells) (Figure 5F, G). On the contrary, cells on stiff gels showed a marked upregulation of TfR-positive EE exocytosis in a larger region around the wound site (Figure 5G, bottom panel). This was especially evident in unrepaired or ‘dying’ cells on stiff gels (Supplementary Figure S7C, D). Most likely, the open wounds in these cells allow sustained Ca^2+^ influx which, in turn, triggers continuous EE exocytosis in regions close to and further away from the wound site to compensate for the membrane resealing defect, albeit without success. In cells on stiff gels that eventually succeed to reseal, the increased membrane tension likely leads to an enhanced exocytosis of EE after wounding which provides the additional membrane required to repair the wounds in these cells (as seen in Figure 5B, bottom panel). The Ca^2+^-mediated EE exocytosis near the wound site potentially lowers membrane tension and thereby facilitates resealing.

The range of EE exocytosis events observed in cells grown on soft and stiff gels yielded the possibility to use these gel experimental setups to study the possible association of wounding-induced CME with changes in membrane tension. Interestingly, laser injury of soft gel-grown HUVEC expressing GFP-Amph1 showed no induction of Amph1-positive endocytosis events after wounding (Figure 5H, I). This suggests that the lowered membrane tension in these cells is adequate for wound repair and that CME is not required to retrieve membrane after resealing and thereby regulate membrane tension variations (note that the wounding-induced EE exocytosis is also minimal in these cells resulting in only a minor increase of PM area) (see Figure 5G, top panel). Examination of CME in HUVEC grown on stiff gels revealed a marked upregulation of Amph1-positive punctae near the wound site after resealing, comparable to the increased EE exocytosis in stiff gels (Figure 5I, bottom panel). Intriguingly, cells on stiff gels that failed to reseal showed hardly any Amph1 positive CME (Supplementary Figure S7E, F) suggesting that EE exocytosis although upregulated probably is not capable of lowering membrane tension to levels that allow resealing and trigger subsequent CME ^[55]^. This supports the notion that Amph1-positive endocytosis events, which are initiated following EE exocytosis, restore membrane tension and homeostasis in wounded cells. Thus, elevated membrane tension as a result of wounding most likely is first relieved by the Ca^2+^-evoked EE exocytosis which is then followed by enhanced endocytosis to restore membrane tension to resting levels at later stages of PM repair.

### Membrane area modulation by exo-and endocytosis is required for tension regulation and wound repair in HUVEC

Given the link between membrane tension and plasma membrane repair we next aimed to assess the role of membrane area modulations in regulating membrane tension for wound repair. As lowering membrane tension by CytoD treatment and cultivation on soft gels permitted efficient membrane repair with only minimal EE exocytosis, we speculated that a general membrane area increase could also lead to similar responses during wound repair. To alter membrane area and assess its role in PM repair, we utilized a recently introduced membrane intercalating agent, azobenzene ^[56]^. This amphiphilic molecule, in its stable *E* – isomer, can intercalate into the plasma membrane of cells and thereby cause membrane area increase. Upon wounding of HUVEC treated with 0.5 mM azobenzene ^[56]^, membrane resealing was unaffected (similar to what was observed with CytoD-treated and soft gel-grown cells) (Figure 6A). Interestingly, HUVEC transfected with TfR-pHuji and treated with azobenzene, showed a marked reduction of EE exocytosis following wounding (Figure 6B, C and Supplementary Figure S8A). As the addition of azobenzene caused minor changes in cell morphology with increased cellular protrusions (see Figure 6B, bottom panel), non-wounding controls of azobenzene-loaded HUVEC were performed revealing that EE exocytosis counts were reduced only after wounding (Supplementary Figure S8B). These results indicate that lowering tension by membrane area increase promotes PM resealing without the need of wounding-induced EE exocytosis.

**Figure 6.**
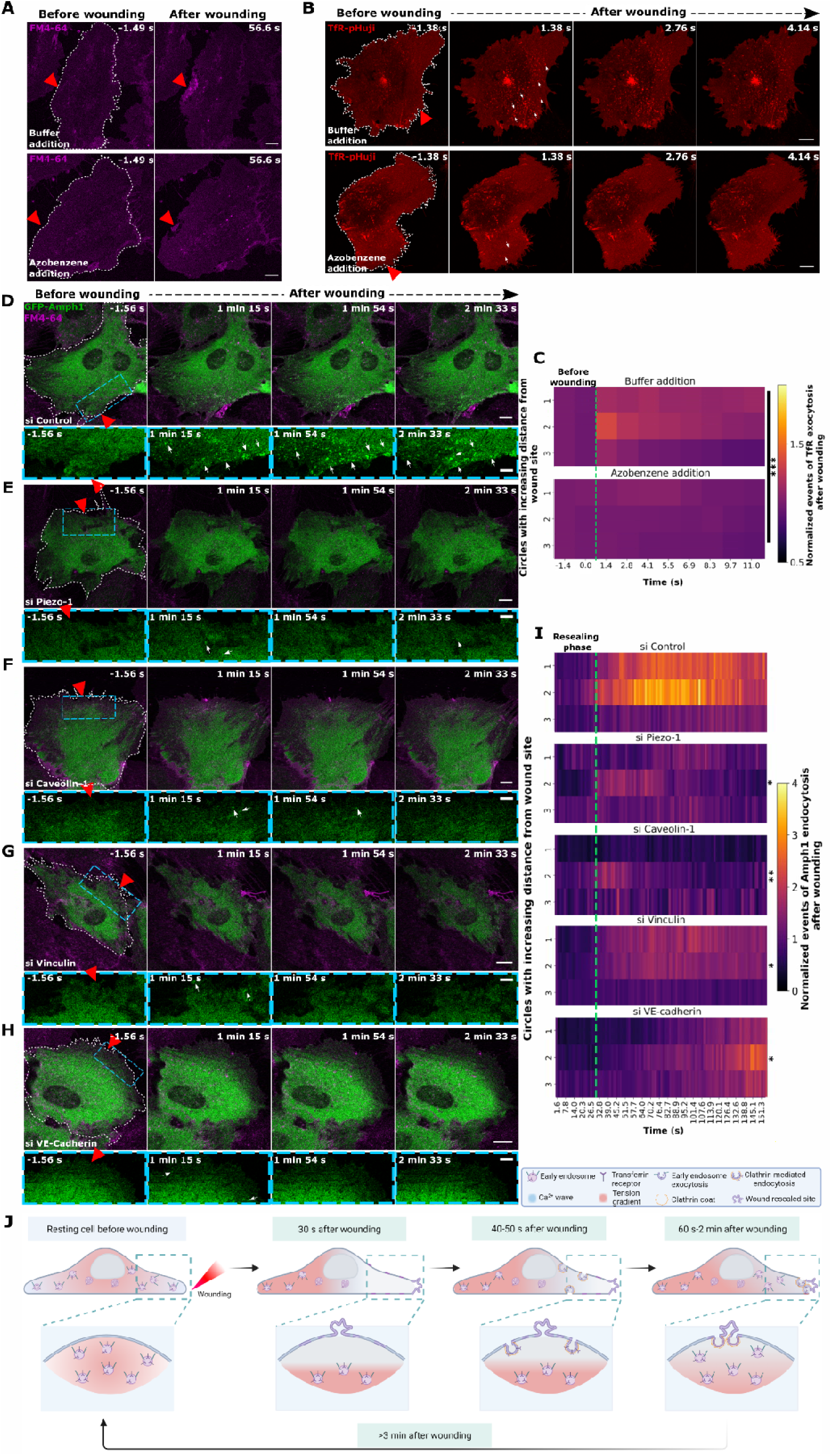
Membrane area-mediated tension changes regulate wound repair. (**A**) HUVEC were treated with the membrane intercalating agent, azobenzene (0.5 mM, bottom panel) or buffer control (top panel), and were laser wounded in the presence of FM4-64 (magenta). Note that efficient resealing occurs in both cases (marked by the FM4-64 delineation site near the wound). (**B and C**) HUVEC expressing TfR-pHuji (red) were laser injured in buffer containing azobenzene (bottom panel) or buffer alone, and assessed for EE exocytosis (**B**). White arrows indicate TfR-pHuji signal increase near the wound indicating EE exocytosis. Quantifications of the EE exocytic events were plotted as a heatmap (**C**) and compared near the wound area (circles 1-3) in cells supplemented with azobenzene or buffer (bottom and top panel, respectively). Plotted events were normalized to the steady-state count before wounding and to the area of the circles used for quantification. The green dotted line signifies the wounding frame. (**D – H**) HUVEC were transfected with control siRNA (**D**) or siRNAs against Piezo-1 (**E**), Caveolin-1 (**F**), Vinculin (**G**), or VE-Cadherin (**H**), together with EGFP-Amphiphysin-1 (green), and wounded in the presence of FM4-64 (magenta). Blue boxed areas below each image depicts a magnification of the area around the wound site for the corresponding EGFP-Amphiphysin-1 signal. White arrows indicate sites of GFP-Amph1-positive CME formed after wounding. (**I**) GFP-Amph1 punctae counts after wounding of the cells transfected with the various siRNA shown in (D – H), were depicted as heatmaps in areas near wound sites (circles 1-3), and after normalization to basal punctae count and circle ROI area. The green dotted line marks the conclusion of the resealing phase after injury. (**J**) Model showing membrane tension changes that regulate EE exocytosis and CME for efficient plasma membrane repair. Red triangles, wound ROI, wounded cells marked by white dashes, and scale bars = 10 μm and 5 μm (zooms). Data are represented as means from 18 - 21 cells for (C) and 21 – 27 cells (I), from 3 biological replicates, and analysed using two-way ANOVA with Tukey’s test. \**P* < 0.05; \*\**P* < 0.01; \*\*\**P* < 0.001.

As the upregulation of CME is only observed after PM wound resealing has occurred, it likely acts to retrieve the membrane added by EE exocytosis and thereby restores the resting membrane tension in the final stages of wound repair. To address this point, we examined whether repair-associated tension regulation by CME is associated with mechanosensitive proteins that had been shown to regulate membrane tension and to associate with various endocytic pathways ^[57]^. We chose four major mechanosensor proteins expressed in HUVEC – Piezo-1, caveolin-1, vinculin, and VE-Cadherin – that were depleted individually by siRNA-mediated knockdown (Supplementary Figure S9A - D) ^[58]^. Wounding of HUVEC with reduced levels of these proteins revealed no major resealing defects (Supplementary Figure 9E), excluding a role of these mechanosensitive proteins in supporting the initial stages of membrane repair, i.e., PM resealing. To assess whether the mechanosensitive proteins are involved in the later stages of PM repair, i.e. the restauration of resting membrane tension by compensatory CME, we analysed GFP-Amph1 positive CME events in cells depleted of the mechanosensitive proteins. Interestingly, a reduction of CME was observed in cells lacking any of the mechanosensitive proteins (Figure 6D – H, I and Supplementary Figure 9F). This suggests that wounding-associated CME is affected by various mechanosensitive proteins to regulate tension in the later stages of wound repair. In summary, wounding-induced EE exocytosis and subsequent clathrin-mediated endocytosis are required to modulate membrane tension and thereby promote resealing and later membrane remodeling, respectively (Figure 6J).

## Discussion

Plasma membrane repair is an emergency response to membrane wounding and involves a repertoire of inherent cellular mechanisms proposed to vary based on cell or wound type ^[11] [12]^. Our results identify a co-ordination of exocytic and endocytic mechanisms that are triggered following wounding and act at different stages to regulate membrane tension changes and thus PM repair. We previously showed that PM wounding induces a Ca^2+^-dependent and spatially restricted EE exocytosis required for membrane area addition to reseal the wound ^[20]^. Endocytosis ^[7] [28]^ has also been associated with wound repair, specifically at the resealing stage. However, we show that CME in HUVEC is only activated after the actual resealing of the wounded membrane and functions at later stages of wound repair. This is in line with the observation that a macropinocytic form of endocytosis occurs after wound repair in cancer cells ^[16]^. Previously, it was also reported that injury-induced exocytosis of lysosomes triggers caveolar endocytosis for the resealing of PM wounds ^[7]^. However, it is difficult to reconcile how both these events mediate the fast kinetics of PM resealing. Rather, we observe a sequential activation of EE exocytosis which provides membrane during resealing, followed by clathrin-mediated endocytosis at a later stage which most likely internalizes the excess membrane deposited and replenishes the pool of EE. This is further substantiated by earlier studies showing that TfR clusters at the PM are classically re-internalized by CME ^[38]^. We also show that the depletion of early endosomes as a result of endocytosis block is detrimental to the initial resealing step, owing to the lack of EE exocytosis and therefore, possibly contributes to the membrane repair defects observed before ^[7] [28]^. A recent study also demonstrated that PM wounds generated by pore-forming toxins as well as laser-induced wounds in various cells initiate EE exocytosis for resealing ^[59]^, validating our previous study ^[20]^, and highlighting that varied wounding sources may not be the predominant factor for initiating different repair mechanisms. Thus, we propose that cells repurpose different cellular mechanisms, EE exocytosis and CME, to mount a spatio-temporally coordinated response to PM injury that regulates membrane tension and thereby aids in efficient PM repair.

Membrane trafficking via exocytosis and endocytosis has been described before to respond to and regulate membrane tension ^[40]^. Specifically, exocytosis reduces PM tension by increasing membrane area and endocytosis decreases membrane area and thus elevates membrane tension ^[41] [60]^. This suggests that the rapid EE exocytosis-mediated increase in membrane area upon wounding and subsequent internalization via CME may be associated with corresponding membrane tension changes. An increased membrane tension at the edges of PM wounds has been long suggested due to the attachment of the PM to cytoskeleton and membrane proteins ^[61]^, and this high tension has been proposed to keep the wound open hindering efficient repair ^[19]^. A previous study reported an association of wounding-induced tension changes with PM resealing ^[19]^, but how this connects to other wound repair mechanisms and the later remodeling steps has not been investigated. Yet other studies revealed an ambiguous role of cytoskeleton-mediated tension changes in PM wound repair ^[62]^. Here, we establish a wounding-induced association of EE exocytosis and CME to sequentially regulate membrane area/tension in the course of wound repair. Following injury, high tension at the wound site is counteracted by the increased membrane area resulting from the immediate Ca^2+^-triggered EE exocytosis that drives resealing ^[20]^. The lowered tension which is a result of this expanded membrane area is then potentially sensed by the cell after the actual PM resealing, possibly via various mechanosensitive proteins such as those analysed here. This initiates compensatory endocytosis eventually restoring membrane tension to resting cell levels, as recapitulated in the experiments performed with cells on gels of varying stiffnesses. The altered membrane tension in these cells, as validated by the tether pulling assays, differs from a recent study which detected no evident changes in membrane tension of fibroblasts and neurons grown on hydrogels of different stiffnesses ^[63]^. However, it is to be noted that the extracellular matrix (ECM) protein laminin used for the adhesion of the cell to the substrate in that study is different from the ECM protein collagen used in our experiments, which may lead to the engagement of different integrins and corresponding tension variations ^[53]^.

Conventional methods of altering membrane tension in cells such as osmotic shocks, stretching, and deadhering ^[42] [55]^, lead to global changes affecting various cellular parameters, possibly including membrane integrity, rendering the methods not suitable for the wound repair studies carried out here. Therefore, we chose to employ more specific ways of modulating membrane tension such as cell growth on gels of different stiffnesses. Actin cytoskeletal rearrangements are also associated with membrane tension changes^[64]^. However, this is distinct from wounding in HUVEC where a direct contribution of the actin cytoskeleton in the initial stages of membrane resealing has not been observed ^[18]^. In summary, we show here that PM wound repair is characterized by membrane tension changes regulated by spatially and temporally coordinated membrane trafficking events - EE exocytosis that occurs immediately after wounding to lower tension for efficient membrane resealing and CME that is activated post resealing and restores the resting membrane tension. Membrane tension is therefore proposed to act as a universal trigger, apart from the well-established extracellular Ca^2+^ influx, of the complex PM repair responses required for restoration of membrane integrity and homeostasis after wounding of cellular membranes.

## Materials and Methods

### HUVEC culture

Primary HUVEC (Human Umbilical cord Vein Endothelial Cells) were acquired from Promocell (C-12203) and cultured in a 1:1 mixture of ECGM-2 (Promocell, C-22111) and M-199 (Pan Biotech, P04-07500), supplemented with serum (10 % FCS for M199 (Capricorn scientific, FBS-11A)), 30 µg/ml gentamicin (Cytogen, 06-03100), 15 ng/ml amphotericin B (Sigma, A2942) and 100 I.U. heparin. Cells were cultured in 37°C, 5 % CO_2_ atmosphere up to 5 passages.

### Plasmids and siRNA

HUVEC were transfected with plasmid DNA or siRNA via electroporation using the Amaxa nucleofection system (Lonza), as described previously ^[65]^. Usually, 0.5 to 5 µg of plasmid DNA was used per 20 cm^2^ of confluent cells.

The following plasmids were purchased from Addgene: TfR-pHuji from David Perrais (# 61505) ^[66]^, Dyn2-pmCherryN1 (# 27689; Christien Merrifield) ^[67]^, PH-PLCD1-GFP (# 51407; Tamas Balla) ^[68]^, mEmerald-Caveolin-C-10 (# 54025) and mEmerald-Caveolin-N-10 (# 54026) were gifts from Michael Davidson ^[69]^. EGFP-Amphiphysin-1 was kindly provided by Harvey McMahon (Cambridge University, UK) ^[70]^ and the following constructs: EGFP-Cdc42, EGFP-Cdc42 N17, EGFP-RhoA, and EGFP-RhoA V14, were generously given by Andreas Pueschel (University of Muenster, Germany). EGFP-CAAX was gifted by Roland Weldich-Soeldner (University of Muenster, Germany). mCherry-CAAX was cloned and provided by Magdalena Mietkowska and Arf6-GFP was generated by Johannes Nass (University of Muenster, Germany). TfR-SEP ^[20]^ and PH-PLCD1-mApple ^[18]^ were described before. Dynamin-2-EGFP was generated in this study by exchanging Dynamin-2 from Dyn2-pmCherry N1 (purchased from Addgene mentioned above) with EGFP from pEGFP-N1 using EcoRI and NheI restriction digestion-based cloning.

For the protein knockdown experiments, 1×10^6^ HUVEC were transfected with the corresponding siRNA (CLTC #1 and CLTC #2 together at 150 nM each, 400nM for Piezo-1, and 600nM each for Caveolin-1, Vinculin or VE-Cadherin) twice for 48 h and the cells were analysed 24 h after the 2nd round of transfection. The two siRNAs against human clathrin heavy chain used in conjunction were purchased from Dharmacon: siGENOME Human CLTC siRNA #1 and #2 (D-004001-01-0004 and D-004001-02-0005, respectively). For the mechanosensor knockdown, the following siRNAs were used: Dharmacon ON-TARGETplus Human PIEZO1 SMARTpool (L-020870-03-0005) against Piezo-1, Dharmacon ON-TARGETplus Human CAV1 SMARTpool (L-003467-00-0005) against Caveolin-1, and Dharmacon ON-TARGETplus Human VCL SMARTpool (L-009288-00-0005) against Vinculin. For VE-Cadherin, the siRNAs used were: Invitrogen Silencer Select siRNA, s223070 (CDH5) and CDH5 synthesized by Microsynth (5′- GGGUUUUUGCAUAAUAAGCTT-3′). As control, non-targeting siRNA AllStars Negative Control siRNA (Qiagen Hilden, 1027281) was used.

### Antibodies and reagents

The following primary antibodies were used: rabbit α-CLTCA (Proteintech, 26523-1-AP, 1:800), rabbit α-Piezo1 (Proteintech, 15939-1-AP, 1:200), mouse α-caveolin1 (BD Biosciences, 610407, 1:1000), rabbit α-vinculin (Proteintech, 26520-1-AP, 1:2000), rabbit α-VE-cadherin (Proteintech, 27956-1-AP, 1:1500), rabbit α -tubulin (Cell Signalling, 2125, 1:2000), mouse α -tubulin (Sigma, T5168, 1:3000), and mouse α-calnexin (BD Trans. Lab, 610524, 1:700). Secondary antibodies employed in Western blots were goat α-rabbit IRDye 800CW (LICOR, 926–32211) and goat α-rabbit IRDye 680CW (LICOR, 926–32221), goat α-mouse IRDye 680CW (LICOR, 926–32220) and goat α-mouse IRDye 800CW (LICOR, 926–32210).

Depolymerization of actin cytoskeleton was induced by Cytochalasin D (Sigma/Merck, C8273) treatment at 1 µM in Tyrode’s buffer (given below) for 30 minutes at 37°C before proceeding to wounding assays. Clathrin- and dynamin-dependent endocytosis was inhibited using Dynasore (Merck Calbiochem, 324410) at 100 µM in Tyrode’s buffer (given below) for 20 min at 37°C before the assays and maintained during imaging. Clathrin-mediated endocytosis was blocked using Pitstop-2 (Abcam, ab120687) at 15 µM in Tyrode’s buffer (given below) for 15 minutes before the imaging assay. Pitstop-2 negative control was used correspondingly in the same experiment (Abcam, ab120688) as a compound control. Fluorescently labelled transferrin uptake (Transferrin-Alexa Fluor 488, Thermo Fisher Scientific, T13342) was also included in the endocytosis inhibitor experiments during the last 5 minutes of treatment time, to estimate the efficiency of clathrin-mediated endocytosis. Membrane area modulation experiments were performed by Azobenzene in the *E*-isomer state at 0.5 mM in buffer added just before imaging ^[56]^.

### Plasma membrane repair assays

### Two-photon laser ablation assay

The two-photon laser ablation assay for membrane wounding was performed as previously detailed ^[23] [20]^. In brief, sub-confluent HUVEC on collagen-coated dishes were imaged in Tyrode’s buffer (140 mM NaCl_2_, 5 mM KCl, 1 mM MgCl_2_, 10 mM glucose, 10 mM HEPES, pH 7.4) (*73*) with either 2.5 mM Ca^2+^ (unless mentioned else) or 100 µM EGTA, and supplemented with 5 μg/ml FM4-64 (Thermo Fisher Scientific, T13320). The ablation and in-situ imaging were performed with an LSM 780 confocal microscope (Zeiss) using a 63× NA 1.4 oil immersion or 40× NA 1.3 oil immersion objective (for the hydrogel experiments) at 37°C. The ablation laser was used at 820 nm wavelength with the power set to 16.5 % (Chameleon Vision NLO laser, Coherent) to generate circular wounds of 2.5 μm^2^ surface area (20 pixels in diameter) on the plasma membrane (image dimensions of 512 × 512 pixels). A time series of 2 frames was recorded prior to ablation to account for bleaching and subsequently, a total of 100 frames were recorded to assess the wound repair dynamics (the frame rate for each experiment is mentioned in the figure timestamps or *x*-axis of the graphs).

After imaging, the time-lapse videos were inspected for cells that showed retraction or *xy* or *z* drifts during imaging and these cells were omitted from the analysis. All analyses were performed on the unmodified image data using Fiji (ImageJ 1.53t) ^[71]^ and custom-written scripts that are available on request. The membrane wound resealing kinetics was analysed using the FM4-64 dye influx with a macro employing the “Plot Z-profile” function of Fiji as detailed in ^[23]^. In short, the fluorescence intensity of the wounded cell was measured for all time frames after wounding. This value for each cell was subtracted from the intensity of a nearby unwounded cell for background signal exclusion, and then normalized to the intensity of the wounded cell prior to injury (frame 1). The subsequent curve from the analysis (in absolute units) indicates successful membrane repair based on a plateau reached by the FM dye increase over time while failure of membrane repair is indicated by a linear increase in the FM dye influx.

### Mechanical scrape assay

Mechanical injury by scraping was performed as outlined before ^[20]^. Shortly, untransfected or siRNA-transfected HUVEC grown to confluence on collagen-coated dishes were treated with the specified inhibitors or proceeded directly for the wounding assay. Cells were scraped in the presence of PBS (+/+ or -/- Ca^2+^ and Mg^2+^ as specified) supplemented with Dextran-Alexa Fluor 488 (200 μg/ml, Thermo Fisher Scientific, D22910) using a cell scraper (Sarstedt, 83.3951) in a gentle and reproducible pattern over the entire dish and allowed to reseal at 37°C for 5 min. The repair process was then stopped with ice-cold propidium iodide (PI) (50 μg/ml, Sigma Aldrich, P4864-10ML) added to the cell suspension for 4 min in ice. Afterwards, the cell suspension was spun down at 200 *g*, 4 min, 4°C, resuspended in cell wash buffer, and analysed by flow cytometry (Guava easyCyte 11 Flow Cytometer, Merck). A total of 5000-10000 cells were measured for each condition, and the fluorescence was analysed using the 488 nm laser line and emission at 525/30 nm for AF488 and 695/50 nm for PI. A strict gating protocol was used to separate the fragmented cell debris and aggregates stained for PI due to the assay nature. Inhibitor treatments generally led to cell loss due to detachments and so, measurements were pooled from multiple technical replicates for each biological experiment. The efficiency of scrape wounding was indicated by the AF488-positive cells (injured) and the PI-positive cells among the AF488-positive cells within the gate of PI signal (injured and non-repaired) indicated the non-resealed cells. The repaired cells were estimated from the fraction of AF488-positive and PI-negative cells within the gated region of PI signal per sample. The efficiency of membrane repair was represented as a percentage of the repaired cells to the total number of scrape-wounded cells.

### Quantification of exocytosis and endocytosis punctae upon wounding

HUVEC were transfected with the EE exocytosis marker (TfR-pHuji or TfR-SEP) or clathrin-mediated endocytosis markers (EGFP-Amphiphysin-1 or Dynamin2-EGFP) and laser wounding was performed as stated above, with or without the various inhibitors used. The analyses of exocytic or endocytic punctae were done as previously described ^[20]^. The events were detected as bright punctae in the corresponding fluorescence channel. In short, pixel classifiers were trained to differentiate between punctae and background using the iLastik software ^[72]^ which utilizes supervised machine learning approaches to generate probability maps. These maps differentiate the signal and background intensities and ensure accurate detection of the exo- and endocytic clusters. The resulting probability maps were thresholded using Fiji and the subsequent analyses were done on the binarized images. An ROI (region of interest)-based analysis was applied on all images to estimate the punctae count (as shown in Supplementary Figure S1G). A set of concentric circular ROIs was generated (each ROI with an increment of 20 µm in diameter from the previous ROI) from the circular wound ROI (2.5 µm^2^), segmenting the wounded cell into circles with increasing distances from the wound site. Circle 1 is closest to the wound site and circle 6 is the farthest from the wound ROI. The number of puncta in these various circle ROIs covering the cell area was measured over time for all the samples. The number of puncta measured at each time point was normalized to the initial number of punctae (before wounding frame) to establish a comparable baseline across cells. Additionally, the puncta count was also normalized to the corresponding ROI area in µm^2^ for all frames to account for the variability in the count due to the differing ROI sizes in different regions of the cell. The null puncta count observed in the initial frame in certain cells was set to 1 to ensure uniform normalization across samples, possibly resulting in an underestimation of the exocytic or endocytic events over time. The increase in EE exocytosis or CME events was plotted over time after wounding in the various circle ROIs as heatmaps using a custom-written Python script (matplotlib, version 3.7.3) ^[73]^. For the punctae count in the endocytic markers (control samples without treatments), a similar analysis was also applied to non-wounded controls where cells were ablated with the laser wounding assay at low laser power (0.2 %). The number of punctae quantified in these cells was used to estimate the baseline for the dynamics of the endosomal markers across *z* and time.

### Nearest-neighbour analysis to assess spatial association of exocytic and endocytic events

HUVEC were cotransfected with TfR-pHuji and the different endocytosis markers (EGFP-Amphiphysin-1 or Dynamin-2-EGFP), and membrane wounding was performed as detailed in the laser ablation assay. The time lapse images obtained were subjected to iLastik pixel classification to generate probability maps of the fluorescence signal. To ensure accurate signal detection and discern the punctate localization of the proteins, probability maps were thresholded and label images were created using scikit image (version 0.21.0) ^[74]^. Punctae were filtered based on size and their counts in the various ROIs were estimated as explained before, and similarly normalized to the initial punctae count and the corresponding ROI area. Next, the TfR-pHuji punctae formed in the first 10 *s* after wounding within circle 1 (region closest to the wound site) were compiled and compared to the endocytic punctae formed across all frames after wounding over the entire cell. This was done using the KDTree algorithm (scikit learn (version 1.3.1), KDTree, metric: *‘minkowski,)* ^[75]^.The distances between the compared punctae were then queried to find the nearest neighbour. For visualization, all TfR-pHuji exocytic punctae (in frames 3 – 10) and their corresponding endocytic punctae over time were mapped on an image (see Supplementary Movie S5). These measurements were also performed with the TfR punctae in circle 6 as a control and for the TfR punctae in circle 2 for EGFP-Amphiphysin-1 (since punctae count analysis shown in Figure 1C detected Amph1 punctae in both circles 1 and 2). Afterwards, the distance of the mapped exocytic and endocytic punctae in the initial frame was used to normalize all the distances measured later to set a baseline for comparison. The normalized mapped distances were then plotted for all the circles. If the punctae of the endocytic protein are spatially associated to TfR exocytosis events, their distances would be smaller and vice versa.

Density plots (seaborn (version ’0.13.0’), kdeplot) ^[76]^ were also generated after the punctae estimation by plotting the distribution of all punctae with respect to normalized distance to the wound site co-ordinates. It is to be noted that a strict thresholding was applied for the EGFP-Amphiphysin-1 and Dynamin-2-EGFP endocytic punctae in these analyses for efficient mapping to the TfR clusters, and this may lead to an underestimation of the number of endocytic punctae in all cells. The nearest-neighbour distance measured in circle 6 acts as a control against randomly distributed punctae of exocytosis and endocytosis that may appear close in the cell. Similar analysis was also performed for non-wounded control cells where random movements of the markers occur over time to thereby estimate baseline association dynamics.

### Hydrogels for tension modulation

Polyacrylamide hydrogels were prepared on Ibidi 2-well glass bottom μ-slides (Ibidi, 80287) following an established protocol with modifications ^[77] [78]^. Briefly, solutions with varying concentrations of acrylamide/bis-acrylamide (3 % / 0.03 % and 8 % / 0.264 %) were mixed to form hydrogels of different stiffnesses (Young’s moduli of 0.2 and 20 kPa, respectively). To allow cell adhesion, the hydrogels were functionalized with a coating of type I collagen using the heterobifunctional linker sulfo-SANPAH. The hydrogel surfaces were covered with a solution of sulfo-SANPAH (Sigma, 1mg/ml, in milli-Q water) and irradiated with 365 nm UV light (intensity of 10 mW/cm^2^) for 1 min. Next, the hydrogels were washed with PBS and incubated with 50 μg/ml rat-tail collagen I (BD Biosciences, 354236) solution in PBS for 2 h at 37°C. After a final PBS wash, the gels were seeded with HUVEC (transfected or non-transfected) for laser wounding experiments. 0.1 x 10^6^ cells were added per 0.2 kPa gel and the cells were imaged 20-21 h post seeding, after ensuring that the cell morphologies looked as expected with less spread cells. For the 20 kPa gels, HUVEC were seeded at 0.14 x 10^6^ and imaged after 22-23 h of growth. The laser ablation experiments were done as previously described using a 40x oil objective, to allow penetration of the two-photon ablation laser light through the gels (∼ 60-80 nm thickness).

Following image acquisition, the time-lapse videos of cells grown on soft gels that showed severe retraction were excluded for future analysis. The efficiency of wound resealing in cells grown on gels was quantified using multiple means. Firstly, the FM4-64 influx over the entire cell was measured as done in the laser ablation assays before (see above). Next, a circle ROI around the wound site of 20 µm in diameter was applied and the intensity of FM4-64 influx in this ROI across time was measured and defined as ‘around wound’ to estimate the delay in membrane resealing seen in a normalized area around the wound site. These intensity measurements were normalized to the whole cell intensity prior to wounding. Finally, the wound delineation sites, marked by the FM-enriched part of the cell after imaging was concluded, were outlined and their areas measured over the different gel conditions. The areas were normalized to the whole cell areas to avoid bias due to the varied sizes of the cells on the different gels (see Figure 5B).

### Tether pulling assay for estimating membrane tension of HUVEC grown on hydrogels

HUVEC were seeded on collagen-coated hydrogels of varying stiffnesses on glass coverslips and cultured as described above. For the tether pulling experiments, 4.5 μm polystyrene particles (Polysciences, 17135-5) were first coated with streptavidin and next with Concanavalin A-biotin conjugate (Sigma Aldrich, C2272) during a 2-hour incubation at 4°C. The biotinylated particles were then suspended in HUVEC medium supplemented with 20 mM HEPES (imaging medium). Just before the tether pulling experiments, a second coverslip was placed on top of the sample, separated by two 400 μm spacers, and the space in between the coverslips was filled with imaging medium containing the biotinylated particles. A droplet of low viscosity mineral oil was used to seal off the inflow area between the coverslips. Tether pulling experiments were performed using a custom-built optical tweezers system, as described in ^[79]^.

In short, the system employed an 808 nm infrared laser (LU0808M250, Lumics GMbH, Germany) with an output of 250 mW. The laser beam was focused onto the sample using a 60× water immersion objective (CFI Plan Apochromat VC 60XC WI NA 1.2, Nikon, Japan) and an objective heater (H401-T-Controller, Okolab, Italy) was used to maintain the sample temperature at 37°C. The back focal plane of the condenser was imaged onto a Position Sensitive Detector (First Sensor, Germany). A cell with sufficient particles nearby was selected for the assay and a single particle was trapped at the focal point of the infrared laser and subsequently brought into contact with the cell at a constant velocity of 1 μm/s. It was held stationary for 1 second to facilitate the binding of Concanavalin A to the cell membrane before being moved away from the cell at a velocity of 0.3 μm/s. The optical force exerted on the particles was then measured by collecting the infrared light. For all the samples, force data were recorded at a rate of 1000 s^-1^, while images were acquired at an approximate rate of 10 s^-1^. This was repeated for various cells across the soft and stiff gels. The force curves were then analysed using a custom Python script. When a tether was formed, the average plateau force was taken as an indicator of the membrane tension.

### Western blotting

Cell lysates were prepared by scrape harvesting HUVEC grown to confluence on a 6 cm dish in cold lysis buffer (50 mM Tris pH 7.4, 150 mM NaCl, 1 mM EDTA, 1% NP-40, 0.5% Na deoxycholate) containing protease inhibitors. Cell suspensions were sonicated for 1 min and centrifuged at 10000 g for 4 min at 4°C. Protein concentrations were determined using the Pierce 660 nm protein assay and after normalization, samples were boiled in Laemmli buffer at 95°C for 10 min. Lysates prepared for Piezo-1 blotting were warmed in Laemmli buffer at 37°C for 10 min. Equal amounts of lysates were subjected to SDS-PAGE and western blotting according to standard protocols ^[80] [81]^. Samples were subjected to 10 % SDS-PAGE at 80 V for 30 min and subsequently at 110 V, and blotted onto 0.2 µm nitrocellulose membranes (Millipore, 10600001) using a wet tank system at 115 V for 1 h at 4°C in Tris-Glycine buffer (25 mM Tris, 190 mM glycine, 20 % (v/v) methanol). The membranes were then blocked in 5% milk in TBST (150 mM NaCl, 50 mM Tris–HCl, 0.1% Tween 20, pH 7.6) for 1 hour and incubated with primary antibodies overnight at 4 °C. For protein signal detection, infrared conjugated secondary antibodies (IRdye 680RD or IRdye 800CW, LICOR) and the Odyssey imaging system (LICOR) were used.

### Statistical analysis

The graphs are represented as means with standard deviations (SD) as error bars, unless indicated otherwise. The number of independent experiments and individual cells from each independent experiment (*n*) are mentioned specifically in the figure legends. At least three biological replicates were performed for each experiment. The analysed data from Fiji were plotted using custom-written Python scripts (Python Software Foundation, version 3.8) or GraphPad Prism 10 (GraphPad Software). Parametric and nonparametric tests were chosen as appropriate for the data based on normality tests and are reported in the figure legends. Specific statistical details for each experiment can be found in the corresponding figure legends. Data were considered statistically significant if the p-value <0.05 (*), <0.01 (**), <0.001 (***), <0.0001 (****) or ns, not significant.

### Data representation

All fluorescence images, immunoblots and time-lapse videos were linearly contrast stretched in Fiji for relevant image representation. All images and movies intended to be compared were processed similarly to avoid any bias. Data were plotted using GraphPad Prism 10.1.0. Figures were assembled for publication using Fiji and Inkscape (San Jose, CA, USA). Animations were created using BioRender.com.

## Data availability

All data are available in the main text or the supplementary information. Additional data or codes related to this paper may be requested from the authors.

## Acknowledgments

We thank Fabian Höglsperger and Bart Jan Ravoo (Center for Soft Nanoscience, University of Münster, Germany) for providing the Azobenzene compound and helpful discussions. We acknowledge Thomas Zobel (University of Münster, Germany) for discussions and support on image analysis. We also thank members of the Institute of Medical Biochemistry and Bioactive Materials Laboratory, for reagents and support. N.R. was a member of CiM-IMPRS, the joint graduate school of the Cells-in-Motion Interfaculty Centre, University of Münster, Germany and the International Max Planck Research School - Molecular Biomedicine, Münster, Germany. This work was supported by grants from the German Research Foundation (CRC1009, project A06, and GE514/6-3 to V.G. as well as BE 6270/3-1 to T.B.), the Max Planck Society (MPG) to B.T., and the CiM Pilot Project programme (PP-2021-01) to N.R. and M.W. B.E.V. and T.B. received funding from the European Research Council (ERC) (PolarizeMe, 771201).

## Author information

N.R. and V.G. conceived the study. N.R. performed the experiments. M.W. and B.T. contributed to the polyacrylamide gel experiments. B.E.V. and T.B. performed the tether pulling assays and T.B. performed mathematical membrane tension calculations. S.W. helped in image analyses planning and developed the codes for analyses. F.B. performed the western blots for the mechanosensor proteins. N.R. and V.G. wrote the original manuscript with inputs from B.E.V., M.W., B.T., T.B., and S.W. All the authors discussed and commented on the manuscript.

## Ethics declarations

## Competing interests

The authors declare that they have no competing interests.

## Notes

### Competing Interest Statement

The authors have declared no competing interest.

