## Supplementary Information for "Membrane tension regulation is required for wound repair"

**This PDF file includes:**

Supplementary Figures. S1 to S9

Captions for Supplementary Movies S1 to S7

**Other Supplementary Materials for this manuscript include the following:**

Movies S1 to S7

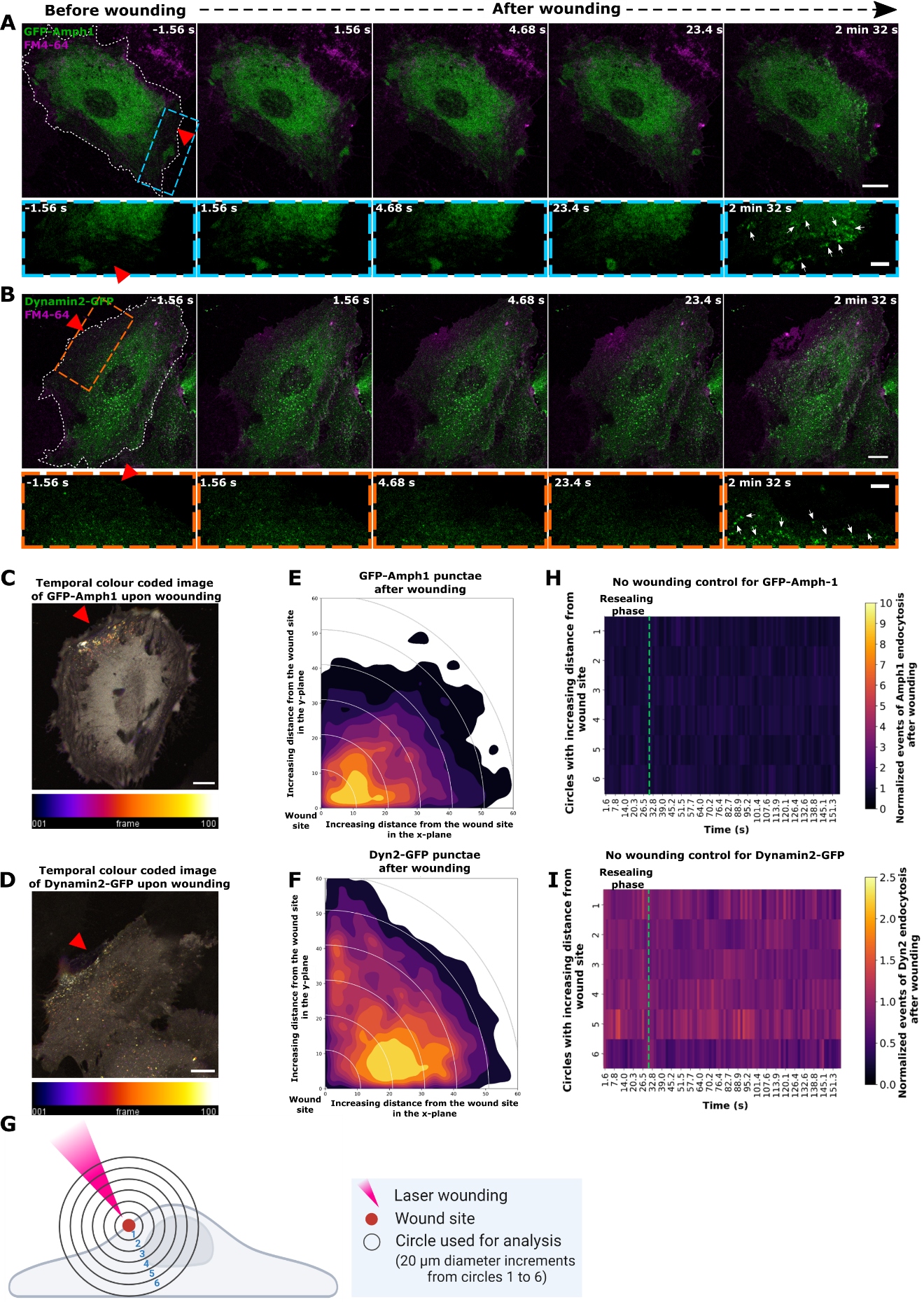

**Figure S1.** **Enhanced clathrin-mediated endocytosis does not occur during wound resealing in HUVEC, but only at later stages of wound repair.** (**A - B**) HUVEC were transfected with markers of clathrin-mediated endocytosis (CME), EGFP-Amphiphysin-1 (labelled as GFP-Amph1) (**A**) or Dynamin-2-EGFP (labelled as Dynamin2-GFP) (**B**) (both displayed in green) and subjected to laser injury in the presence of the membrane-impermeable wounding dye, FM4-64 (magenta). Representative images of the same cell before and after wounding are shown here to detect the resealing response and corresponding dynamics of the endocytic proteins. Boxed areas below show higher magnifications around the wound site at each time point for EGFP-Amphiphysin-1 (blue box) and Dynamin-2-EGFP (orange box). Note that successful resealing was observed around 20-30 *s* after wounding, as detected by the containment of the FM4-64 dye influx around the site of wounding (time point 4 for both panels). No changes in GFP-Amph1 (A) or Dynamin-2-EGFP (B) were observed during this time, but a significant enhancement of both markers as punctate structures occurred at a later time point after resealing (last time point for both panels). Red triangles, wound sites. White dashes, wounded cells. White arrows indicate the endocytic punctae formed around the wound site, which are absent in the previous frames. Scale bars, 10 μm and for zooms, 5 μm. (**C and D**) Temporally colour-coded images of CME events upon wounding of HUVEC expressing GFP-Amph1 (**C**) or Dynamin2-GFP (**D**) (corresponding to images in Figure 1A and B, respectively). The images are generated from all time frames and show the endocytic punctae formation with respect to time after wounding, represented in the colour scale below each image (with frame 1 = -1.56 s and frame 100 corresponding to 2 min 33 s after wounding). Apart from the restricted localization of CME punctae around the wound site, note that the punctae close to the wound are brighter coloured indicating that they are formed in the later stages of repair. Red triangles, wound sites and scale bars, 10 μm. (**E and F**) Density plot representations of GFP-Amph1 punctae (**E**) or Dynamin2-GFP punctae (**F**) formed in HUVEC after wounding. Here, the distribution of the punctae from all time frames is plotted based on the distance to the wound site, with the wound co-ordinates being set to (0, 0) (origin of the graph). Note that the majority of the punctae formed by both endocytic proteins are close to the wound site, suggesting a wounding-induced response and not steady-state endocytosis. Punctae counts in the frames before wounding are also included here, which possibly explains the widespread distribution of Dynamin-2 structures owing to its punctate pre-wounding localization all over the cell unlike cytosolic GFP-Amph1. (**G**) Scheme showing the quantification strategy used for endocytic punctae count during wound repair ^[20]^. The wounded cell is segmented into 6 concentric circles with increasing distance from the wound site (each circle with increments of 20 μm in diameter from the previous circle). Circle 1 is closest to the wound site while circle 6 is the farthest from the wound region. Endocytic punctae are counted in each circle area that is separated from the other circles, and are normalized to the punctae count in the frame before wounding as well as the circle area (due to the varying sizes of the circles). Wound site, red circle on the cell membrane also highlighted by the red triangle (indicating laser ablation site). (**H and I**) HUVEC were transfected with EGFP-Amphiphysin-1 or Dynamin-2-EGFP and subjected to laser injury with a low laser power, used as a non-wounding control. In the absence of membrane damage, the endocytic punctae positive for EGFP-Amphiphysin-1 (**H**) or Dynamin-2-EGFP (**I**) were quantified (as in Figure 1, C and D) and represented as heatmaps here. The endocytic punctae are plotted as in the quantification scheme shown in (G) with respect to time after wounding. The green dotted line indicates the time of resealing generally observed in wounded cells. Note that no significant increase in endocytic punctae is seen over time ruling out major changes in steady-state endocytosis. Means are plotted for (E), (F), (H) and (I) from 18 – 25 cells compiled from 3 biological experiments. Statistical comparisons were performed to estimate the effect of wounding on endocytic punctae count in different regions of the cell over time by using one-way ANOVA with Kruskal-Wallis test and obtained *P* = 0.1229 for (H) and *P* = 0.1141 (I).

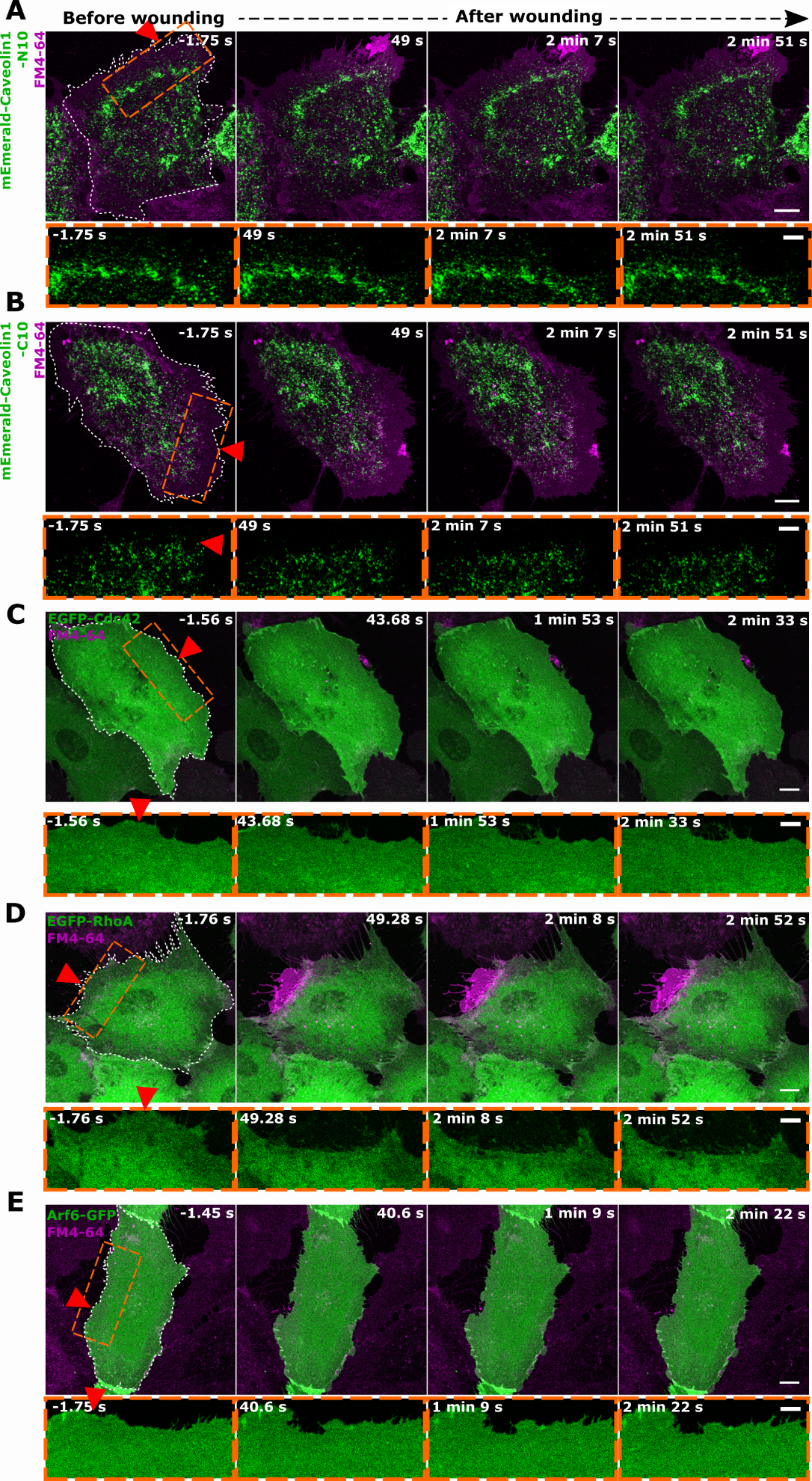

**Figure S2.** **Clathrin-independent endocytosis is not upregulated at the wound site during HUVEC PM repair. (A - E)** HUVEC were transfected with the following markers of clathrin-independent endocytic pathways: mEmerald-Caveolin-N-10 (**A**), mEmerald-Caveolin-C-10 (**B**), EGFP-Cdc42 (**C**), EGFP-RhoA (**D**), or Arf6-GFP (**E**) (all displayed in green), and subjected to laser injury in the presence of the membrane-impermeable dye, FM4-64 (magenta). Representative images of the same cell before and after wounding are shown here to assess the dynamics of clathrin-independent endocytosis during membrane wound repair.

The wound site is highlighted by the orange box and the corresponding magnifications are shown below for each fusion protein over time. Observe the lack of induction of endocytic punctae positive for these marker proteins around the wound site, unlike observed with the markers for clathrin-mediated endocytosis (amphiphysin-1 and dynamin-2) indicating a specific association of CME with HUVEC PM repair. Caveolin-1 localization was tracked using fluorescent reporter constructs tagged at the N- or C-terminus of the protein, in accordance with the reported overexpression artefacts observed in other cells ^[70]^ (even though no such phenotype was observed in HUVEC after transfections of these constructs). Red triangles, wound sites and white dashes, wounded cells. Scale bars, 10 μm and magnifications, 5 μm.

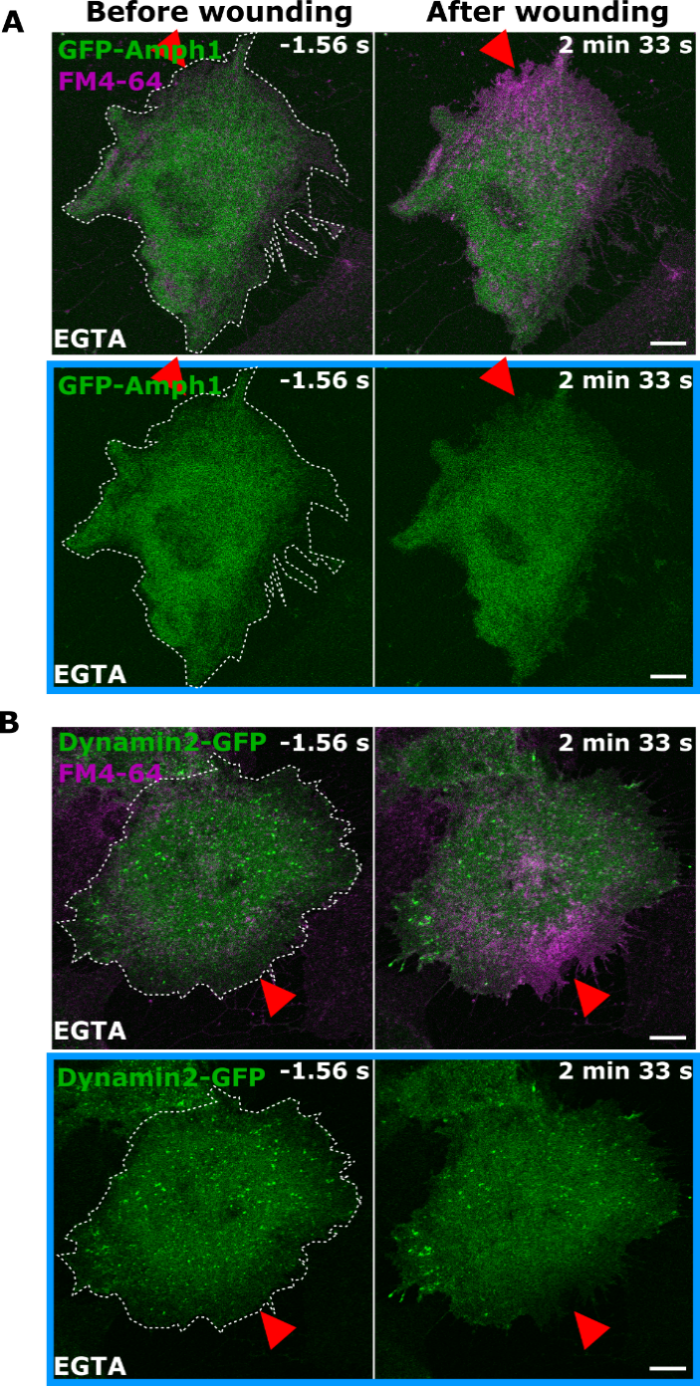

**Figure S3.** **Enhanced clathrin-mediated endocytosis is dependent on Ca^2+^-triggered HUVEC membrane resealing.** (**A - B**) HUVEC expressing CME markers, EGFP-Amphiphysin-1 (**A**) or Dynamin-2-EGFP (**B**) were subjected to laser injury in buffer containing the Ca^2+^ chelator, EGTA and FM4-64 dye (magenta and everything else displayed in green). Representative images before and after wounding are shown here. Impaired membrane resealing in the presence of EGTA is revealed by the increase in cellular FM4-64 intensity. No change in the localization of the CME proteins can be observed around the wound site when membrane resealing is impaired in the presence of EGTA. The GFP channels alone are highlighted below each panel in blue boxes. Red triangle, wound site and white dashes, wounded cells. Images representative of 3 – 4 independent experiments. Scale bars, 10 μm.

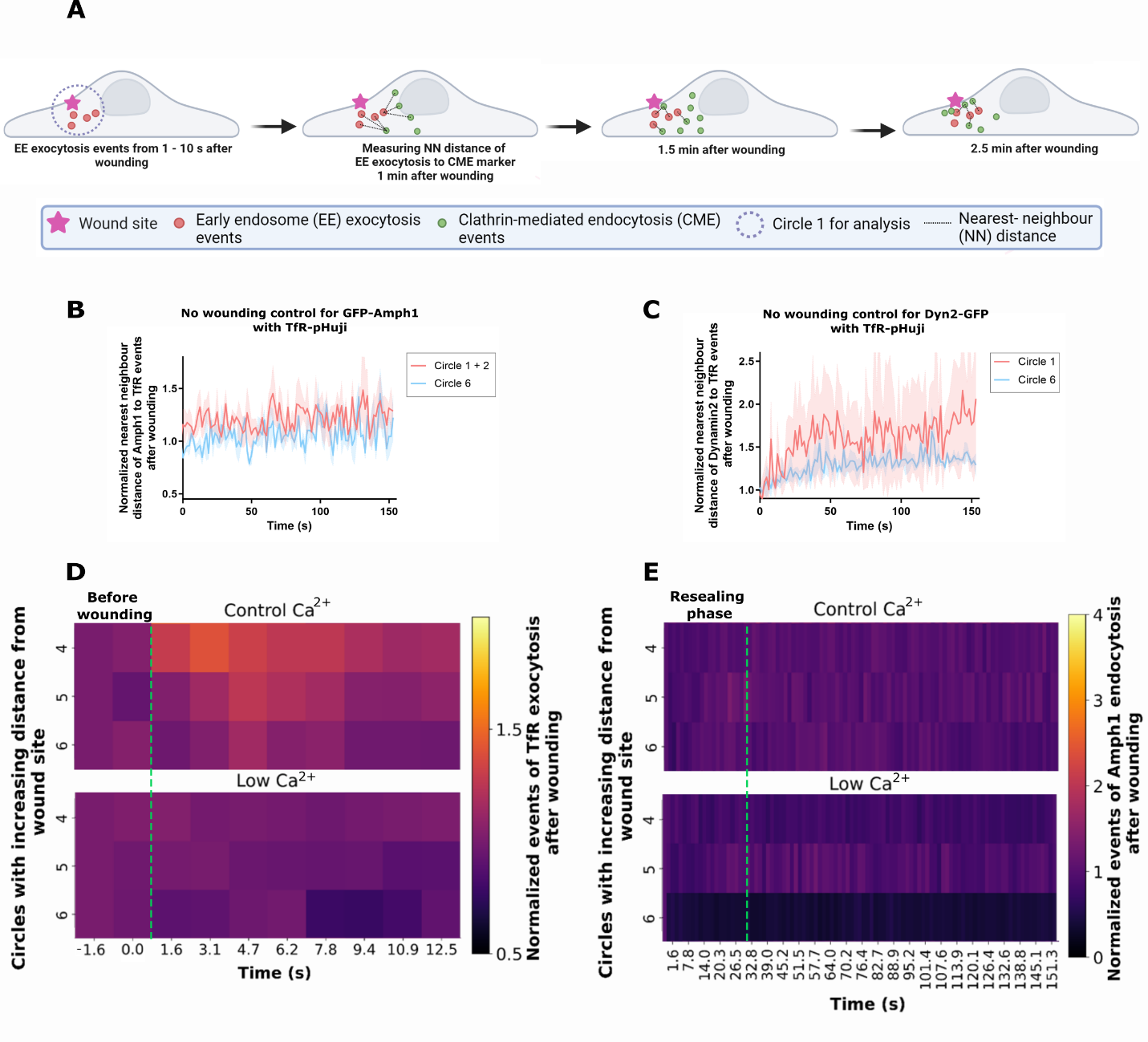

**Figure S4.** **Nearest neighbor analysis of exocytotic and endocytic events to assess their spatial association upon membrane wounding. (A)** Schematic showing the nearest-neighbour analysis used to measure the spatial association of EE exocytosis and CME during wound repair. HUVEC transfected with EE exocytosis marker, TfR-pHuji and CME marker (EGFP-Amphiphysin-1 or Dynamin-GFP) were laser injured and time-lapse videos were recorded (as in Figure 2A and B). An image of the TfR-pHuji clusters (appearance of bright fluorescence due to pH neutralization after exocytotic fusion) near the wound site (circle 1) in the first 10 *s* after wounding was generated to map all EE exocytosis events close to the wound site (marked as red punctae in the animation). The distance of these TfR clusters to all endocytic punctae (shown as green punctae) formed over the entire course of repair (up to 2 min 33 s after wounding) was measured, averaged across various clusters, and calculated as the ‘nearest-neighbour’ distance (represented in grey lines). If the endocytic punctae formed near the EE exocytosis events, the nearest neighbor distance would be lower and vice versa. The nearest-neighbour distance across all time frames was normalized to the distance before wounding (also see Supplementary Movie S5 and Materials and methods for further details). Note that depending on the localization and kinetics of the endocytic protein analysed, the initial distance may vary. (**B and C**) Nearest-neighbour analysis of low laser wounding controls for TfR-pHuji with EGFP-Amphiphysin-1 (**B**) or with Dynamin-2-EGFP (**C**). Note that the curves for circles near and far away from wound sites do not vary for both endocytic proteins over time. This excludes a role of stochastic association between EE exocytosis and CME events in contributing to the association seen during wound repair. Mean ± SEM with error bars as the fill area around each curve. n=18 - 20 cells over 3 independent replicates. (**D and E**) Quantifications of punctae count following wounding of TfR-SEP (**D**) and GFP-Amph1 (**E**) expressing cells in the presence of low Ca^2+^ (bottom panel) or control (2.5 mM Ca^2+^, top panel) are plotted as heatmaps. Only the regions far away from wound site (circles 4 – 6, also see Supplementary Figure S1G) are shown here for both conditions (corresponding to the other circles shown in Figure 2, I and K). The green dotted line in the heatmap indicates the time point of wounding (F) shown to identify EE exocytosis occurring immediately after wounding and resealing phase occurring around 30 *s* after wounding in (G) to delineate the endocytic punctae formed in the later stages of repair. Means from 24 – 31 cells over 3 independent replicates. Statistical analysis using two-tailed Mann-Whitney U test was performed for (B) and (C) with *P* = 0.4233 and *P* = 0.0596 respectively, and two-way ANOVA with Tukey's test was done with *P* = 0.0610 for (D) and *P* = 0.0234 (E).

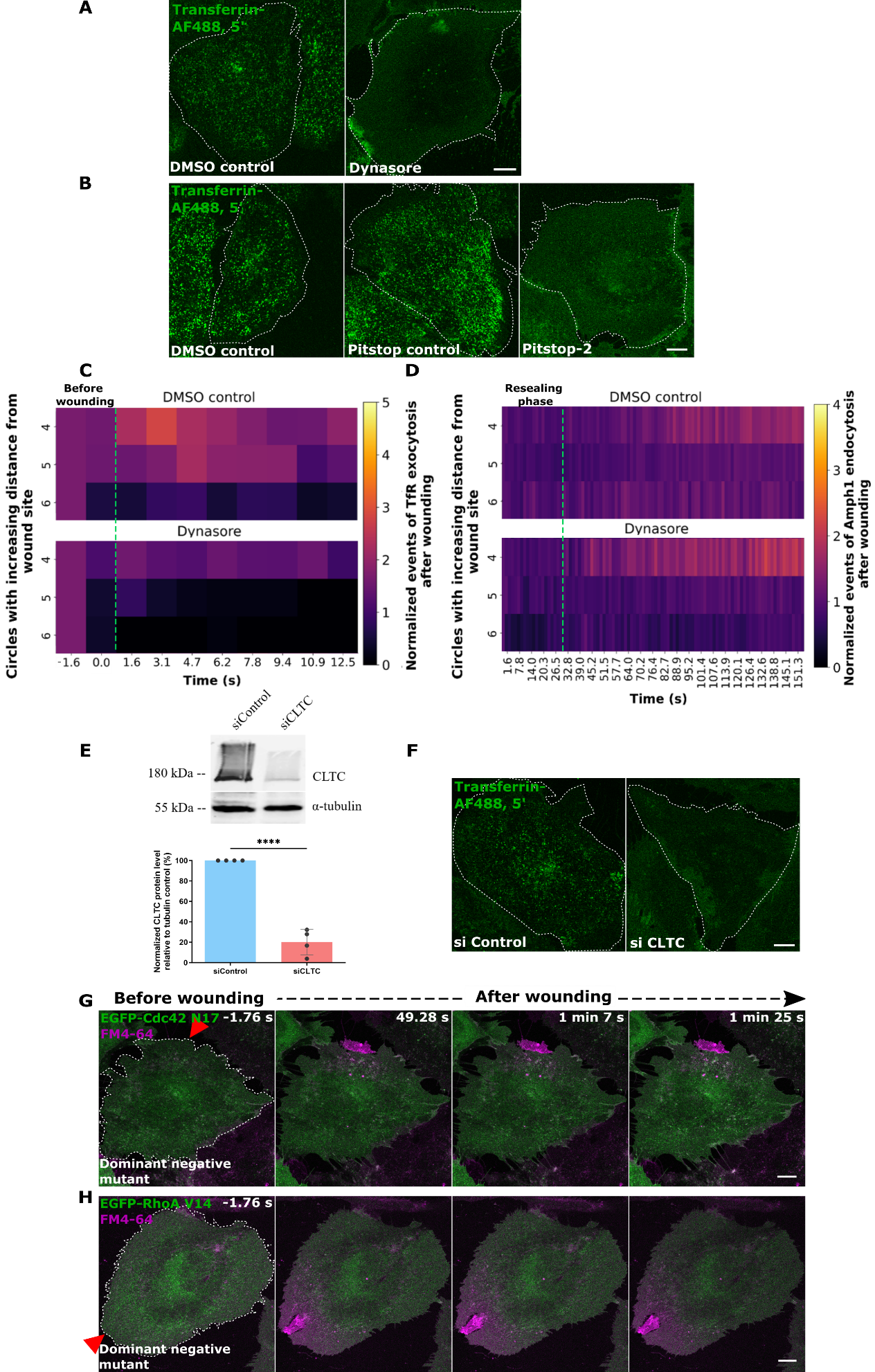

**Figure S5.** **Inhibition of clathrin-mediated and clathrin-independent endocytosis in HUVEC shows varied effects on wound repair.** (**A and B**) HUVEC were treated with DMSO (left panels) and CME inhibitors, Dynasore (**A**) or Pitstop-2 (**B**) and the efficiency of endocytic uptake was demonstrated by a short transferrin (Tf) pulse (Transferrin-AF488) for 5 min (shown in green). As a negative control for Pitstop-2, the cells were also treated with Pitstop control, which has a similar structure to Pitstop-2 but does not block CME. Representative images of Tf uptake into early endosomes are displayed here. Note the lack of internalized transferrin signal indicating impaired dynamin- and clathrin-mediated endocytosis. (**C and D**) Regions far away from the wound site (circles 4 – 6) were quantified over time after Dynasore treatment for: exocytosis marked by TfR-SEP (**C**) and endocytic events labelled by EGFP-Amphiphysin-1 (**D**) and plotted as heatmaps (related to the circles 1-3 shown in Figure 3, G and H). Exo- and endocytic punctae counts are normalized for each circle ROI and to the initial events before wounding. The green dotted lines in the heatmaps label the wounding time point (C) and the end of resealing phase in (D). (**E**) Immunoblot showing levels of clathrin heavy chain (CLTC) after siRNA transfection with siControl or pooled siCLTC (top panel) and represented as a percentage of the loading control, α-tubulin here (bottom panel). (**F**) HUVEC were transfected with siControl or pooled siCLTC and the inhibition of endocytosis was assayed as in (A) and (B), by a transferrin pulse for 5 min (shown in green). Note that the siCLTC samples showed an inhibition of endocytic uptake and thus lack of Tf-positive early endosomes. (**G and H**) HUVEC were transfected with the dominant negative mutant of Cdc42, GFP-Cdc42 N17 (**G**, green in image) or the dominant negative mutant of RhoA, EGFP-RhoA V14 (**H**, shown in green as well), and laser wounded in the presence of FM4-64 dye (magenta). The dominant negative isoform of Cdc42 is known to specifically inhibit the clathrin-independent CG pathway ^[30]^ while the RhoA V14 isoform inhibits RhoA-dependent endocytic pathways ^[83]^, aiding to evaluate the functional role of these endocytic pathways in HUVEC membrane repair. Representative images after wounding show no defects in HUVEC resealing as revealed by the confined localization of FM4-64 to the wound site, thus excluding roles for CG and RhoA-dependent pathways in HUVEC PM repair.

Red triangle, wound ROI and white dashes outline wounded cells. Scale bars, 10 μm. Means plotted for (C) and (D) and mean ± SD plotted for (E), pooled from 3 - 4 independent experiments. Statistical analysis was performed with two–way ANOVA with Tukey’s test with *P* = 0.2161 for (C) and *P* = 0.6102 (D), and with unpaired two-tailed Student’s *t*-test for (E). *****P* < 0.0001.

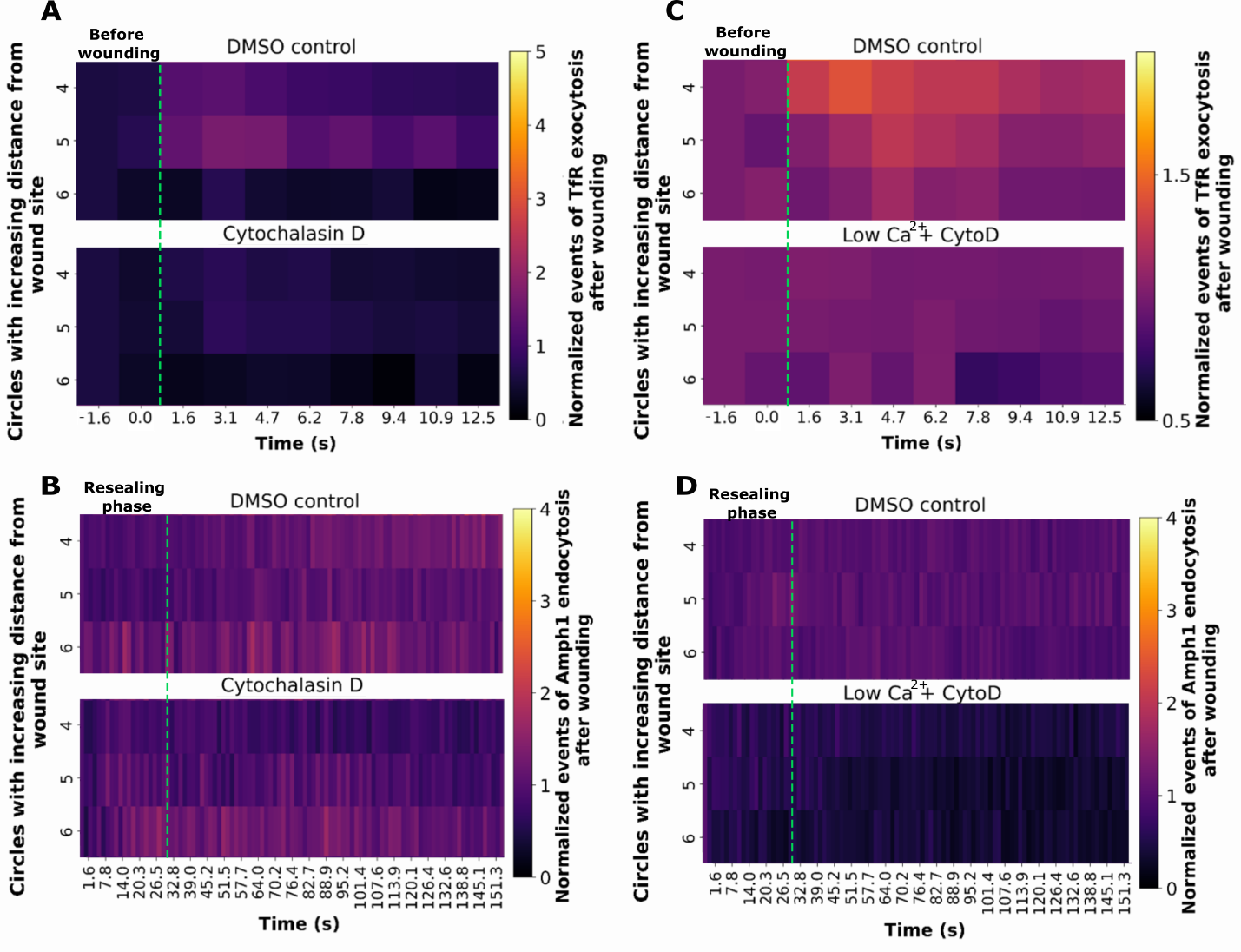

**Figure S6.** **Tension variations do not affect EE exocytosis and clathrin-mediated endocytosis in regions far away from the wound site.** (**A and B**) Punctae analysis of EE exocytosis events positive for TfR-SEP (**A**) and CME events marked by EGFP-Amphiphysin-1 (**B**) was performed after wounding in DMSO control (top panel of each analysis) or Cytochalasin D treated HUVEC. The results are plotted as heatmaps over time for circles far away from the wound site (circles 4 - 6) and shown here after normalization to counts prior to wounding and circle area (refer to Figures 4F and G for circles 1-3). The green dotted line indicates the time of wounding for the EE exocytosis count map (A) and the time point of resealing for CME count representation (B). (**C and D**) HUVEC treated with low Ca^2+^ and Cytochalasin D (indicated as CytoD) were wounded and assessed for TfR-SEP count (**C**) or GFP-Amph1 count (**D**), as in (A) and (B). The heatmaps show regions far away from the wound site, revealing no major changes in exo- and endocytosis following treatments (see Figures 4K and M, for circles 1-3).

Means are plotted for all heatmaps pooled from 21 – 31 cells from 3 independent experiments. P values calculated using two-way ANOVA with Tukey’s test showed *P* = 0.0149 for (A), *P* = 0.0856 (B), *P* = 0.0679 (C) and *P* = 0.056 (D).

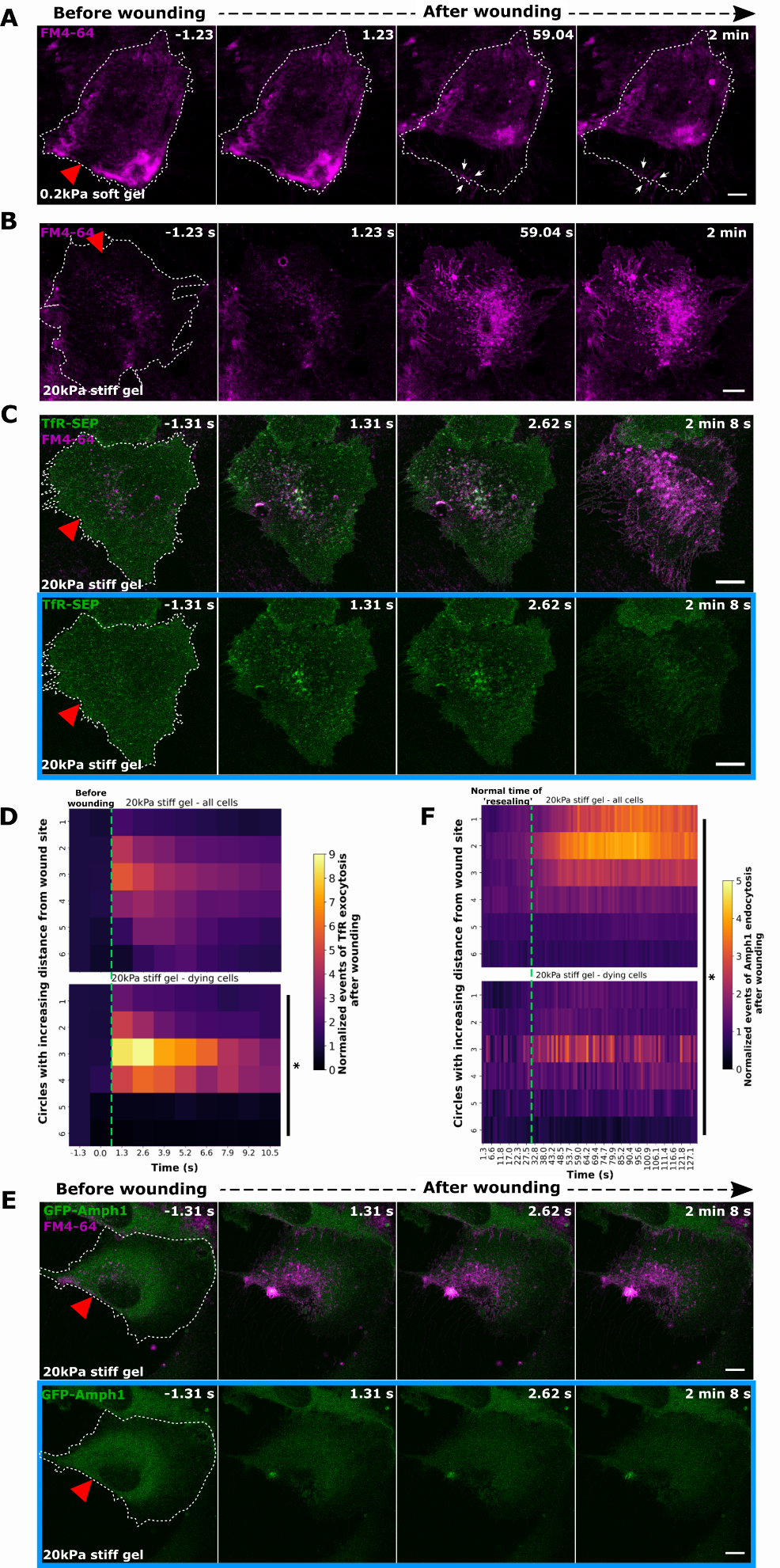

**Figure S7.** **High membrane tension is associated with compromised membrane repair and reduced EE exocytosis and CME around the wound site in HUVEC (A)** HUVEC grown on 0.2 kPa soft gels were laser injured in the presence of extracellular FM4-64 dye (magenta). Still images before and after wounding are shown here. Note the retraction of the entire cell following wounding revealed by the FM4-64 dye labelling of the plasma membrane footprint. This phenotype is observed in 30-40 % of cells grown on soft gels. The white dashes outlining the wounded cell over the time frames after wounding show the footprint of the cell. Also note that the FM4-64 dye influx is still restricted around the wound site (highlighted with white arrows in the image), albeit the lack of attachment of the cell, indicating successful membrane repair in these cells. (**B**) HUVEC cultured on 20 kPa stiff gels were wounded in the presence of FM4-64 dye (magenta) added to the buffer and images before and immediately after wounding are shown. Massive FM4-64 influx into the entire cell indicates defective membrane resealing. Also observe the FM4-64 accumulation at the wound site in the frame after wounding (similar to Figure 4A, bottom panel). Membrane repair was defective in 30-40 % of wounded cells grown on stiff gels. **(C)** HUVEC grown on 20 kPa stiff gels were transfected with TfR-SEP (green) and laser injured in the presence of wounding dye, FM4-64 (magenta). Images after wounding show compromised membrane repair as indicated by the continued influx of FM4-64 dye throughout the cell. Boxed areas below in blue display the GFP channel only (TfR-SEP) for each time point. Note the enhanced EE exocytosis punctae (as indicated by TfR-SEP fluorescence increase) all over the cell. This is probably owed to the open membrane wound in these dying cells which permits sustained Ca^2+^ influx and initiates excess EE exocytosis. (**D**) Quantifications of TfR-SEP exocytosis events after wounding are plotted as heatmaps for all cells grown in 20 kPa stiff gels (top panel) and only for non-repaired cells on 20 kPa stiff gels (bottom panel). Data are normalized and plotted similar to Figure 5G. EE exocytosis close to the wound site is comparable in both populations of stiff-gel cells while the exocytosis events in regions distant from the wound site are upregulated in the dying cells, correlating to a prolonged Ca^2+^ influx through the open wound and a response to deal with the membrane-repair defects observed. **(E and F)** HUVEC were transfected with EGFP-Amphiphysin-1 (green), grown on 20 kPa stiff gels and laser injured in the presence of FM4-64 dye (magenta). Representative still images of a cell showing repair defect after wounding are displayed in (**E**). The Amph1-GFP channel is highlighted below for all the time panels in the blue box. Note the lack of Amph1-endocytic events as opposed to those seen in repairing-competent cells grown on stiff gels, indicating an absence of compensatory endocytosis in non-repaired cells. This is quantified and plotted as heatmaps over time across the entire cell in (**F**) and compared to Amph1 events in all cells on stiff gels (top panel). No significant Amph-1 endocytic events were observable for dying cells grown on stiff gels, suggesting that high tension is detrimental for membrane repair and later restoration of membrane homeostasis.

Red triangle, wound ROI, white outline, wounded cells, and scale bars, 10 μm. Means plotted for all graphs from 3 independent experiments with *n* = 12 – 33 cells (D) and 7 – 27 cells (E) (the cell count for the dying cells corresponds to 30 – 40% of the total stiff gel-grown cells). Statistical comparisons were performed as follows: one-way ANOVA with Kruskal Wallis test to compare punctae count across various regions of the cell in (D, bottom panel) with *P* = 0.0203 and in (F, bottom panel) with *P* = 0.4046, and two-way ANOVA with Tukey’s test to compare punctae count in all cells on stiff gels and dying cells on stiff gels with *P* = 0.4664 (D) and *P* = 0.0334 (F).

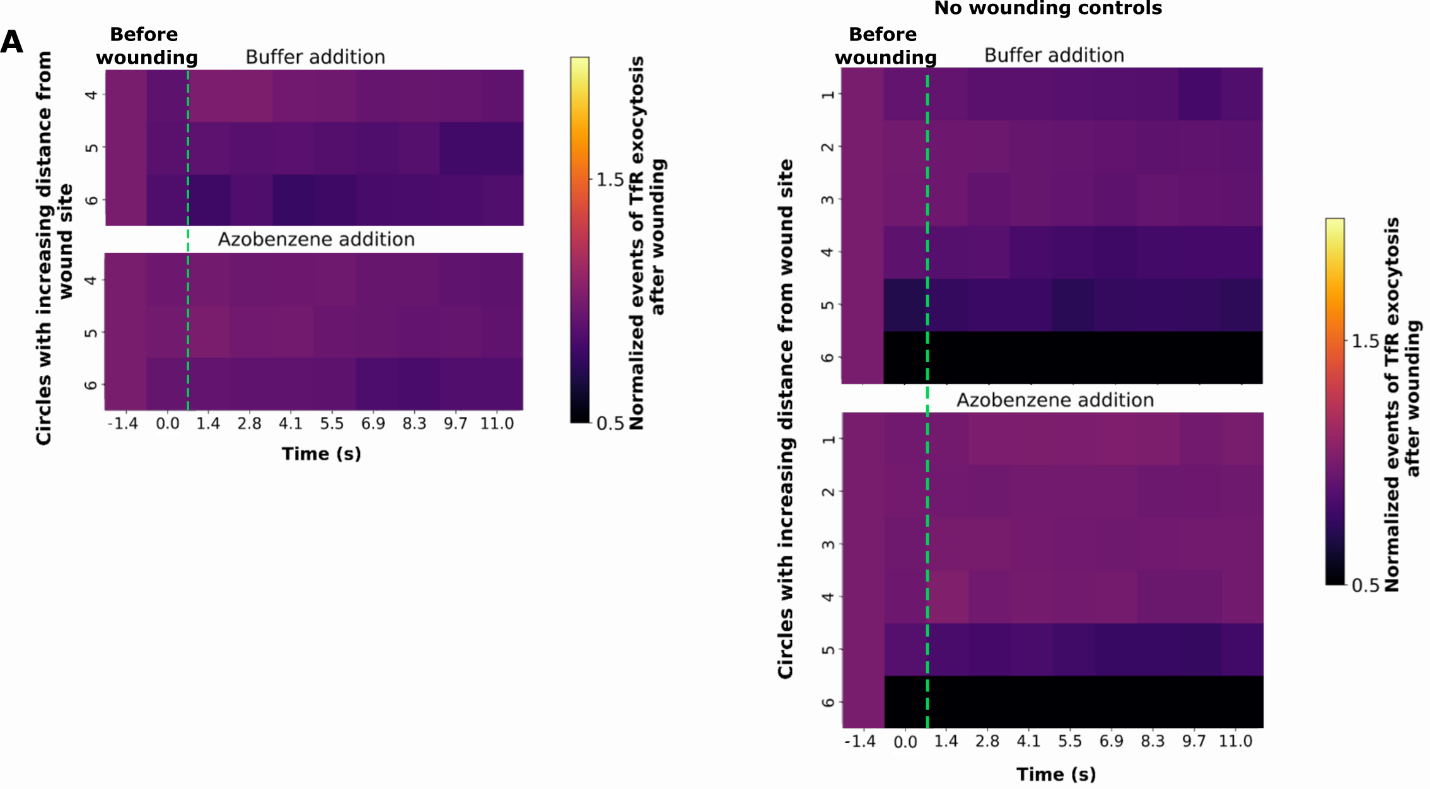

**Figure S8.** **Membrane area increase by azobenzene does not affect EE exocytosis in distant parts of wounded cells or steady-state exocytosis in unwounded cells.** (**A)** Punctae analysis of EE exocytosis events positive for TfR-pHuji was carried out after membrane injury of HUVEC treated with control (buffer addition, top panel) or azobenzene addition (bottom panel). Data are plotted for regions far away from the wound site (circles 4 – 6) over time after normalizations (corresponding graph for circles 1-3 in Figure 6C). (**B**) HUVEC kept in buffer alone or azobenzene containing buffer were wounded at a low laser power (non-wounding controls) and TfR-pHuji positive EE exocytosis counts were quantified as in (A) and plotted for the entire cell (circles 1 – 6), shown here as heatmaps. No measurable EE excoytotic events were noted in contrast to the areas around the wound sites in wounded cells (Figure 6C). This excludes an effect of mechanical stimulation that may occur during exchange of solutions (buffer or azobenzene containing buffer) or off-target effects of azobenzene in affecting basal exocytotic processes in cells (without wounding). The green dotted lines in both heatmaps indicates the time of wounding.

Means plotted here from 18 – 21 cells for (A) and (B). Statistical comparisons were performed using two-way ANOVA with Tukey’s test with *P* = 0.2264 for (A) and *P* = 0.4857 for (B).

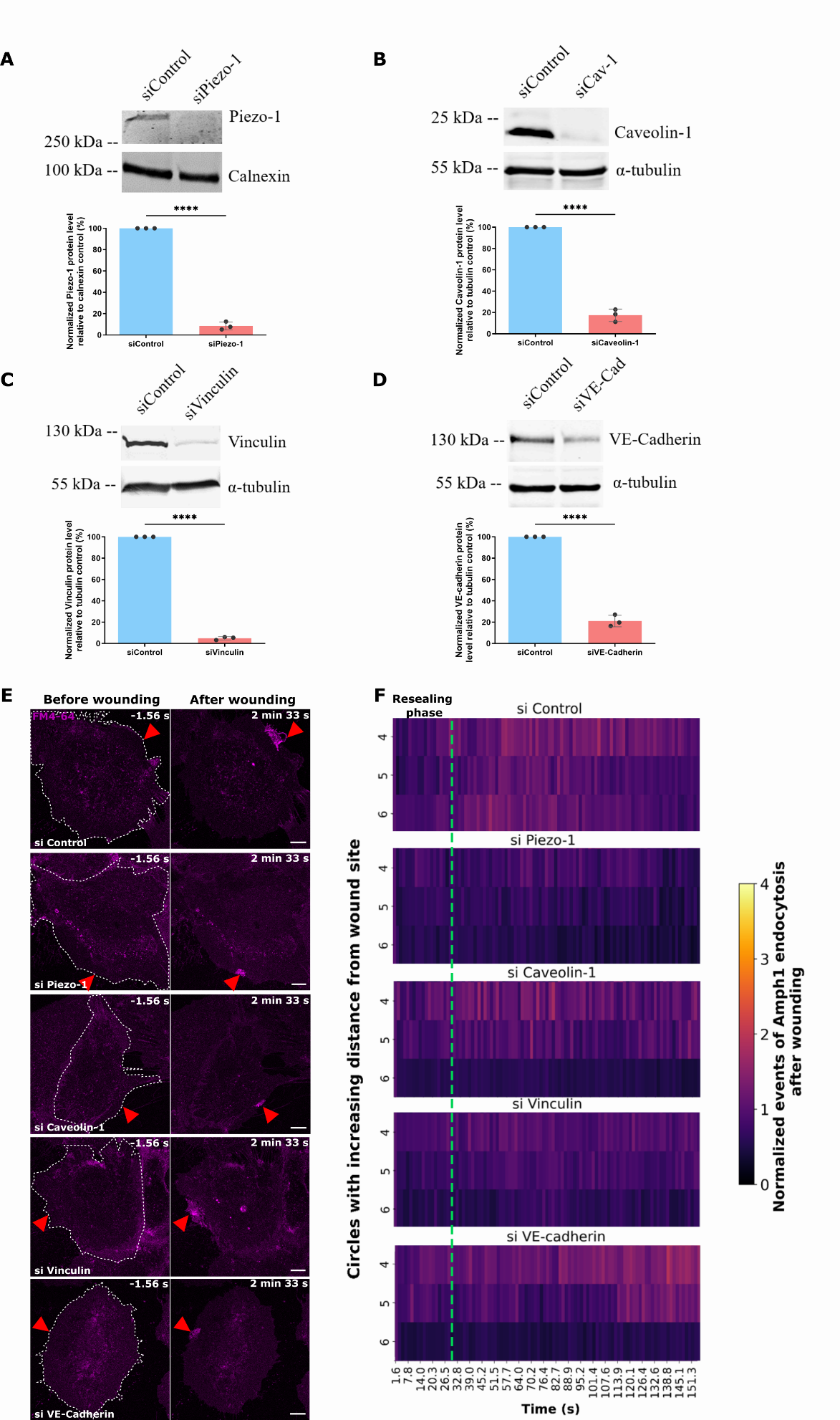

**Figure S9.** **Depletion of different mechanosensor proteins in HUVEC does not affect membrane resealing but inhibits endocytosis at later stages of membrane repair.** (**A - D**) HUVEC were transfected with siControl or siRNAs against different mechanosensitive proteins and knockdown efficiencies were assessed by Western blots of total lysates probed with the respective antibodies directed against Piezo-1 (**A**), Caveolin-1 (**B**), Vinculin (**C**) and VE-Cadherin (**D**). Calnexin (A) or α – tubulin (B – D) were used as loading controls (top panels). The graph at the bottom of each panel shows the analysis of the knockdown efficiency of each siRNA, plotted as a percentage normalized to the loading control. Significant depletion of all mechanosensitive proteins was observed. (**E**) HUVEC membrane resealing was analysed after depletion of the different proteins shown in (A-D). Following siRNA transfection, cells were laser injured in the presence of FM4-64 (magenta). No major defect in membrane resealing was observed in all cases, indicated by the delineation of the FM4-64 dye influx to the ablation site similar to the control sample. Red triangles, ablation sites. White dashes, cell outlines. Scale bars, 10 μm. (**F**) HUVEC transfected with siRNAs targeting the different mechanosensor proteins (as shown in E) and co-transfected with EGFP-Amphiphysin-1 were analysed for CME events after wounding. Punctae counts in regions far away from the wound site (circles 4 – 6) are plotted as heatmaps (circles 1-3 shown in Figure 6I). The green dotted line indicates the completion of the resealing phase. No marked change in Amph1 endocytic punctae was observed for all samples in varying parts of the cells.

Mean ± SD plotted for 3 independent experiments for (A – D) and means plotted for (F) from 21 – 27 cells combined from 3 biological replicates. Unpaired two-tailed Student’s t-test was performed for (A – D) with *P* < 0.0001. For the comparisons of punctae count of each siRNA transfection with siControls in (F), two-way ANOVA with Tukey’s test was utilized with *P* = 0.0823 (Piezo-1), 0.2765 (Caveolin-1), 0.2028 (Vinculin), and 0.5822 (VE-Cadherin).

**Movie legends**

**Movie S1.**

Time lapse recording of laser ablation of HUVEC shows the appearance of CME punctae (EGFP-Amphiphysin-1, green) after membrane resealing (FM4-64 wounding dye in magenta), related to Figure 1A. Red triangle, wound ROI. White arrows indicate Amph1 punctae that appear after wounding in the later stages of repair. Frames were captured at 1.56 *s* intervals. Time is displayed in minutes and t = 0 *s* represents the time of wounding. Scale bar, 10 μm.

**Movie S2.**

Time lapse recording of laser ablation of HUVEC shows the appearance of CME punctae (Dynamin-2-EGFP, green) after membrane resealing (FM4-64 wounding dye in magenta), related to Figure 1B. Red triangle, wound ROI. White arrows indicate Dynamin-2 punctae that appear after wounding in the later stages of repair. Frames were captured at 1.56 *s* intervals. Time is displayed in minutes and t = 0 *s* represents the time of wounding. Scale bar, 10 μm.

**Movie S3.**

Time lapse recording of laser ablation of HUVEC shows the appearance of clusters of cell surface transferrin receptor (TfR-pHuji, red, increased fluorescence after exocytosis-induced pH neutralization) immediately after membrane wounding and the subsequent appearance of CME (EGFP-Amphiphysin-1, green) in the later stages of repair (related to Figure 2A). Red triangle, wound ROI. White arrows indicate the spatial association of Amph1 CME events with the TfR clusters post wounding. Images were taken every 1.49 *s*. Scale bar, 10 μm.

**Movie S4.**

Time lapse recording of laser ablation of HUVEC shows the appearance of transferrin receptor (TfR-pHuji, red) clusters immediately after membrane wounding and the subsequent appearance of CME (Dynamin-2-EGFP, green) in the later stages of repair (related to Figure 2A). Red triangle, wound ROI. White arrows indicate the spatial association of Dynamin2 CME events with the TfR clusters post wounding. Images were taken every 1.49 *s*. Scale bar, 10 μm.

**Movie S5.**

Video showing an example of the nearest neighbor analysis used to measure the spatial association of EE exocytosis events near the wound site (in circle 1) (TfR-pHuji segmented punctae, red) and clathrin-mediated endocytosis (EGFP-Amphiphysin-1 segmented punctae, green) after membrane wounding. Red triangle, wound ROI. Circle 1 is marked by the white concentric circle covering the cell. Blue lines indicate the distances measured between the EE excoytotic punctae in circle 1 and all the Amph1 punctae. Video taken with a frame interval of 1.56 *s* is shown for the TfR punctae combined from the first 10 *s* of wounding and the nearest-neighbour analysis for Amph1 is shown for the entire time series. Scale bar, 10 μm.

**Movie S6.**

Time lapse recording of laser ablation of HUVEC grown on 0.2kPa soft gels shows efficient membrane resealing (FM4-64 dye in magenta) (related to Figure 5B). Red triangle, wound ROI. White arrows indicate the delineation of the wound resealing site in the later stages of repair. Frames were captured at 1.31 *s* intervals. Time is displayed in minutes and t = 0 *s* represents the time of wounding. Scale bar, 10 μm.

**Movie S7A.**

Time lapse recording of laser ablation of HUVEC grown on 20 kPa stiff gels shows delayed membrane resealing (FM4-64 dye in magenta) (related to Figure 5B). Red triangle, wound ROI. White arrows indicate the large wound resealing site indicating the enhanced FM dye influx and delayed repair. Frames were captured at 1.31 *s* intervals. Time is displayed in minutes and t = 0 *s* represents the time of wounding. Scale bar, 10 μm.

**Movie S7B.**

Time lapse recording of laser ablation of HUVEC grown on 20 kPa stiff gels shows inhibited membrane resealing (FM4-64 dye in magenta) (related to Supplementary Figure S7B). Red triangle, wound ROI. Frames were captured at 1.31 *s* intervals. Time is displayed in minutes and t = 0 *s* represents the time of wounding. Scale bar, 10 μm.
